# How many phage species remain undiscovered? Species sampling approaches to inform phage discovery

**DOI:** 10.64898/2026.02.15.704868

**Authors:** Massimo Cavallaro, Andrew Kinsella, Spyridon Megremis, Andrew Morozov, Andrew D. Millard, Fabian Freund

## Abstract

The emergence of antimicrobial resistant bacteria has been identified as one of the most serious public health and development threats for the near future. The use of bacteriophages (phages) is a promising solution for the sustainable control of these pathogens. Phages are natural viral predators of bacterial pathogens. However, due to the variability and adaptability of bacteria, developing effective and sustainable phage treatments requires drawing from a wide variety of different phage species. This study applies specialised mathematical and computational estimation approaches to the problem of sampling and discovering species of phages in microbiological communities. Our goal is to predict how many new phage species will be discovered in future samples based on species recorded in a phage sequence database for eight common bacterial hosts. While internal validation shows that current estimators yield low prediction errors for how many new species can be found when sampling more phages, these critically assume that the underlying mechanisms of phage sampling and database inclusion remain similar across time - an assumption put into question by our analysis for all but the host genus *Mycobacterium*.

From this, we discuss how our results can still inform future sampling strategies - both through directly predicting the number of additional species, or by detecting changes in sampling and database inclusion.

Our results have the potential to inform and optimise the hunt for and isolation of novel of phages from the the natural environment.

**Highlights:**

- Species abundance distribution of phages infecting seven of eight analysed host bacterial genera collected in phage databases changed significantly over time.
- Maintaining current sampling strategies for phages of host *Mycobacteriut* may lead to diminishing returns in the number of new phages found in additional samples.
- Both non-parametric and parametric estimators of the numbers of unseen species show comparable, moderate errors when predicting a random subset from the remainder of hostspecific phage sets from the INPHARED database

## 1. Introduction

Bacteriophages, or phages, are viruses that specifically target and kill bacteria. Phages are the most abundant biological entities on our planet, their estimated number is *10*^*31*^ particles. Phages comprise an extremely vast and diverse group of bacterial viruses that do not share a common evolutionary origin [1]. They are characterised by their common ability to parasitise, prey on, and destroy bacterial cells [2]. Phages play key role in controlling microbial ecosystems and regulating bacterial populations, as they act as natural predators in every environment, including humans. Phages have been explored as an alternative to antibiotics since the 1920s, particularly for combating multi-drug-resistant bacterial infections [3]. This is related to the currently growing antimicrobial resistance (AMR), the ability of microorganisms (mainly bacteria) to survive antibiotic treatments. AMR is considered to be a global health threat, largely driven by the overuse and misuse of conventional antibiotics. In humans, AMR contributes to increased mortality rates and prolonged hospital stays, making infections harder and more expensive to treat [4]. In agriculture, development of AMR results in selecting and spreading resistant genes and bacteria across food chains [5].

Despite being highly specialised bacterial predators, phages are not immune to resistance mechanisms put in place by bacteria to defend themselves. Bacteria can evolve strategies to evade phage attacks, which may reduce the effectiveness of phage therapy over time and result in the same resistance associated with traditional antibiotics [6]. One promising approach to counter this phenomenon and provide sustainable control of pathogens is the use of so-called phage cocktails, which are combinations of different phages tailored to target a specific bacterial infection [7]. This strategy mimics natural ecosystems, where a single bacterial species is often targeted by a wide diversity of phage species [8].

To efficiently implement phage therapy, using targeted cocktails, there is a strong need for maintaining extensive libraries of characterised phages. Scientific efforts are currently underway to discover and catalogue new phages (e.g., the phage database INPHARED [9]), and the findings suggest that the diversity of phages is too far from being fully explored. This leads, however, to the fundamental open question: How many phage species remain undiscovered? A related, but more relevant practical question is: How many new phage species should we expect to observe if we sample additional *m* phages?

In this paper, based on the curated collection of phage genomes from INPHARED, we explore the biodiversity of phages observed in eight common bacterial pathogen host genera (i.e., *Escherichia, Klebsiella, Mycobacterium,
Pseudomonas, Salmonella, Staphylococcus, Streptococcus*, and *Vibrio*). Using the advanced mathematical techniques borrowed from the species sampling problem literature, we estimate the number of yet-to-be-discovered species in future samples. We assess different parametric and non-parametric estimators for the number of additional phage species in further samples and show that most of them perform moderately and comparably well. We also discuss and analyse the critical assumption of these approaches that the species abundances entering INPHARED are following a similar distribution over time, an assumption we can show is likely violated for most host genera. We then discuss how this prediction approach still can provide informative insight to improve future repeated sampling of phages from the same environment.

## 2. Methods

Mathematically, the question that we want to answer is: how many new phage species can we expect in additional samples based on the existing data. We address this question in the context of the phage diversity in humans. We should note that this question is part of the broad ‘species sampling’ problem (SSP), which uses a variety of general inference and estimation approaches in statistics [10]. We apply and compare predictions from several existing estimation methods borrowed from the SSP theory to a well-curated phage database.

### 2.1. Data source

Phage genomes and their metadata were retrieved from the September 2024 and the May 2025 versions of the INPHARED database [9], hereafter referred to as DB24 and DB25, respectively. Phage genomes were clustered into species using previously described method [11], where a species was defined as 95% identity over 85% target coverage. The bacterial host was resolved at the genus level in the database.

### 2.2. Notation

For each bacterial host we assume that, from a full population of *N* individual phages hosted, a random sample of *n, n* ≪ *N*, individuals *(X*_*1*_, …, *X*_*n*_) is observed and recorded. These are drawn from the same underlying species set/abundance distribution. Each individual phage also belongs to one of *S* species, with *S*_*obs*_*(n)* being the number of species actually observed in the sample. We are mainly concerned with the observed abundance *a*_*i*_*(n)* of individuals of species *i, i = 1*, …, *S*_*obs*_*(n)*, and the numbers *f*_*k*_*(n) :=* [{*i*[*a*_*i*_*(n) = k*}[of species observed *k* times, *k = 1, 2*, …, with 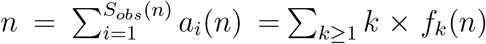 and *S*_*obs*_ *= Σ*_*k*≥1_ *f*_*k*_*(n)*. The true (unknown) species relative abundance distribution, underlying 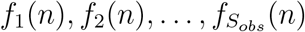, does not depend on *n* and is denoted by *p*_*0*_, *p*_*1*_, *p*_*2*_, …, *p*_*S*_, and the observed frequency count follows a multinomial distribution with total number of event *S*_*obs*_ and event probabilities *p*_*k*_*/(1* − *p*_*0*_*), k = 1, 2*, …, *S*.

### 2.3. The species sampling problem - estimators for unseen species

As the sampling effort increases, and so the number of observed individuals, it is expected that previously unobserved phage species in the initial sample will be observed in the new sample. The additional sample *(X*_*n+1*_, …, *X*_*m*_) is drawn from the same underlying species set/abundance distribution as the already observed sample *(X*_*1*_, …, *X*_*n*_) The estimation of the number of novel species *u(n, m)* in a new sample of size *m* and ultimately of the total number of unseen species *S*_*obs*_*(m + n) = S*_*obs*_*(n) + u(n, m)* is a central component of the SSP, which describes statistical questions around estimating and inferring properties of the underlying distribution of species based on limited data from that distribution. See [12] for an in-depth discussion of SSPs.

We estimated *u(n, m)* using two classes of estimators, non-parametric and parametric. We consider three non-parametric estimators. Two are based on the Good-Toulmin estimator [13] (GT)

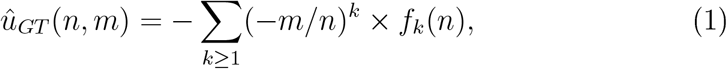

which is referred to as non-parametric since it does not rely on any assumption regarding the distribution *p*_*k*_ underlying *f*_*k*_*(n)*. For *m* ≥ *n*, Σ_*k*≥1_(−*m/n)*^*k*^ in Equation (1) does not converge (gives large weights to high abundance classes), and thus the Good-Toulmin estimator tends to oscillate strongly in this case, leading to suboptimal estimation properties. Several correction terms (smoothings) have been proposed to dampen these oscillations. One correction was proposed by Efron and Thisted,

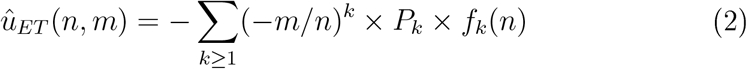

with 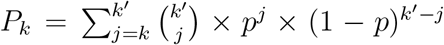[14], where *p = 1/(1* − *m/n)* and *k*^′^ is a tunable parameter, which we choose to be *k*^′^ *= 10*. We will refer to this estimator by the abbreviation ET. In [10] it was shown that this can be seen as averaging over truncations of the sum (1) at a random, binomially distributed summand index. Additionally, the study also suggests a more optimal (binomial) truncation distribution for *m > n*, see [10, Table 1]. Technically, this just changes *(P*_*k*_)_*k*∈ℕ_. We will refer to the estimator using this alternative smoothing as OSW. Note here that we use smoothing in ET for all *m*, while only for *m > n* for OSW (for *m* ≤ *n*, OSW is GT). A further non-parametric estimator was proposed in [15], which only uses the count of species appearing a single time or twice in the sample. This estimator will be referred to as CJ and is defined as

**Table 1:** Observed numbers of individual phages (*n*) and phage species (*S*_*obs*_) per bacterial host in the 2024 and 2025 datasets.

| Host | 2024 data |  | 2025 data |  |
| --- | --- | --- | --- | --- |
| | $n$ | $S_{obs}$ | $n$ | $S_{obs}$ |
| *Escherichia* | 2559 | 716 | 2863 | 826 |
| *Klebsiella* | 1502 | 532 | 1821 | 616 |
| *Mycobacterium* | 3123 | 563 | 3195 | 574 |
| *Pseudomonas* | 1644 | 448 | 1878 | 517 |
| *Salmonella* | 1384 | 332 | 1584 | 370 |
| *Vibrio* | 1089 | 476 | 2381 | 602 |
| *Staphylococcus* | 912 | 279 | 1005 | 313 |
| *Streptococcus* | 938 | 449 | 1331 | 705 |

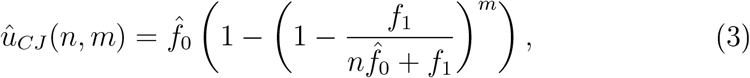

where 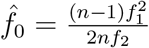 if *f*_2_ > 0 and 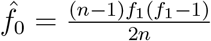 if *f*_2_ > 0.

The second type of estimators is derived from the assumption that the species abundances *p*_*k*_ follows a distribution taken from a parametric family of distributions. The first family consists of Poisson-gamma mixtures (also often referred to as a negative binomial distributions) for *k = 0, 1, 2*, …, i.e.,

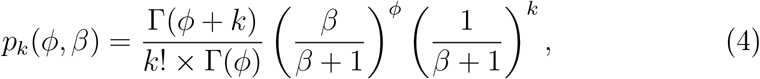

where *Γ* is the Gamma function and the parameters *ϕ* and *β* are to be learned from data. The expected number of unseen species corresponds to the mass at *k = 0* and, under the Poisson-gamma assumption, reads

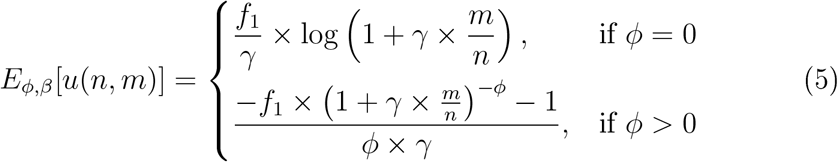

with *γ = 1/(1 + β)* [14, 16]. We obtained maximum likelihood estimates 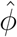 and 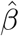 of the parameters *ϕ* and *β* from the observed frequencies *f*_*k*_ by maximising the multinomial likelihood function

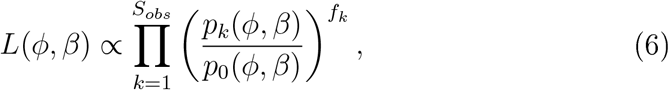

which yields the estimator

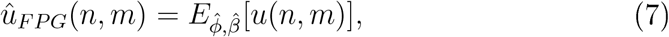

hereafter referred to by the abbreviation FPG (Fisher-Poisson-gamma).

Our second parametric distribution family is given by the Pitman-Yor two-parameter model. The Pitman-Yor model is a class of general distributions of set partitions for integer sets {*1*, …, *n*} with a flexible structure to describe increasing the set sizes sequentially [17] and has been proposed as a fairly general underlying distribution family for species sampling problems [18]. In more detail, the probability *P* of observing counts 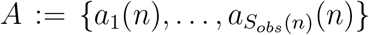 is assumed to depend on the two parameters *(α, θ)* via

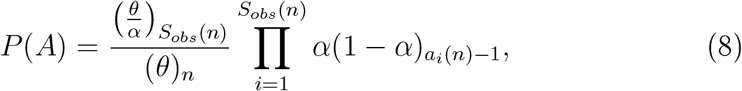

where 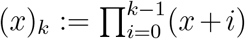 denotes the rising factorial. Following [12], we use an empirical Bayes (plug-in) estimator where we first provide the maximum likelihood estimate 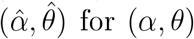 for *(α, θ)*. Then, we can estimate the number of unseen species as the expected value of *u(n, m)* under the corresponding distribution, which has a closed expression for given *(α, θ)* [12],

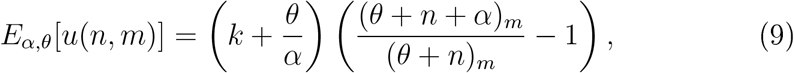

thus leading to the Pitman-Yor-prior (PYP) estimator

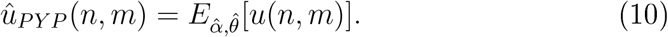

For PYP and FPG, we computed the maximum likelihood estimate via numerical optimisation using the Nelder-Mead optimisation method as implemented in the R package `optimx` [19, 20]. Both parametric estimators allow to capture the observed species diversity in general, though not perfectly - justifying their use as estimators in our use case. Supplementary Figures 2 and 3 show the fit of the ML parameter estimate to the species abundance spectrum.

A critical assumption of all these estimation methods is that both the already observed sample of size *n* and the additional sample of size *m* are sampled from a common underlying (unknown) species (abundance) distribution (e.g., [10]). As the phage research community sequentially enters more entries into the database, this stationarity is not necessarily given for future samples as sampling strategies, sampling targets (e.g. specific host species or environments) and/or which sequences are sent to the database may change over time. We will assess and address this issue below.

Moreover, we can only make any predictions on the distribution of phage species entering the database. This is not the true ecological phage species abundance distribution. However, for potential clinical use the isolated phages, from which database entries are coming, are indeed the relevant distribution.

### 2.4. Validation

To assess how well the estimators OSW, ET, CJ, FPG and PYP perform in the specific setting of species sampling of bacteriophages, we used a data-driven approach of internal validation. For each bacterial host, we used random subsets of a given size (corresponding to 25%, 35%, 50%, 60% and 80% of the sampled phages, rounded to the next integer) to predict the new species in remainder of the full set from this host (i.e., minus the random subset). We then measure the normalised absolute error (NAE), i.e. 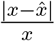 for a true value *x* compared to its estimate 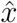. We performed 500 iterations of this validation approach. We performed this analysis on the two temporal snapshots DB24 and DB25 of the database.

For the two estimators with overall best performance from the internal validation (OSW and FPG), we further assessed how prediction accuracies compare if we fix the training set size and predict the number of additional species for different sizes of sets of newly sampled phages. We used training set sizes of 200, 300, 400, 500 and new sample sizes *m* = 100, 200, 300, 400, 500 to perform predictions from the training sets (and then compare with a real subset of the same size). Both the training and the prediction sets were random (disjoint) subsets of the database at the snapshot from 2025. Again, we performed 500 validation runs.

A known issue with the species sampling problem is that all estimates depend on how representative the observed sample is for the underlying species abundance distribution, which is hard to assess, see e.g. [21, 22]. At least for assessing temporal representativeness, i.e. whether the underlying species distribution of the recorded species changes over time, we can assess this as follows: We applied the different estimators, based on input from DB24, to predict the number of additional phage species per host genus in DB25. We compared the error of this estimation (NAE) with the error of predicting, from a random subset of DB25 of the same size as the number of phage species of this host in DB24, the number of new phage species in the rest of the DB25 snapshot.

### 2.5. Prediction from latest snapshot

We projected the estimated number of new species in *m* newly sampled (isolated) phages, based on the DB25 data set, using the recommended OSW estimator for each host, as well as using FPG and CJ. As the OSW estimator is not guaranteed to be monotone in *m*, we also reported its monotone- concave modification following Appendix 6 in [10]. The modification is 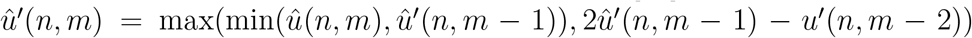, with 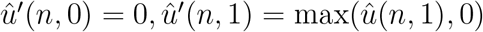.

Confidence intervals were obtained by means of bootstrap at the species distribution level (see Supplementary Text 1 for more details).

## 3. Results

### 3.1. Descriptive analysis

The 2024 data set contains 29,410 records of individual phages spanning 10,934 species, observed across 309 bacterial host genera. The 2025 data set includes 36,288 individual phages belonging to 14,604 different species from 318 bacterial host genera. For the eight bacterial hosts considered, i.e., *Escherichia, Klebsiella, Mycobacterium, Pseudomonas, Salmonella, Staphylococcus, Streptococcus*, and *Vibrio*, the total numbers of observed individual phages *n* and observed phages species *S*_*obs*_ are reported in Table 1. The distributions of the number of observed species counts *f*_*k*_ are illustrated in Supplementary Figure 1 and Figure 1 for the 2024 and 2025 data sets, respectively. They all exhibit fat tails, with many rare phage species observed one or two times, and a few species observed many times, sparsely populating the tails of the empirical distributions.

**Figure 1:**
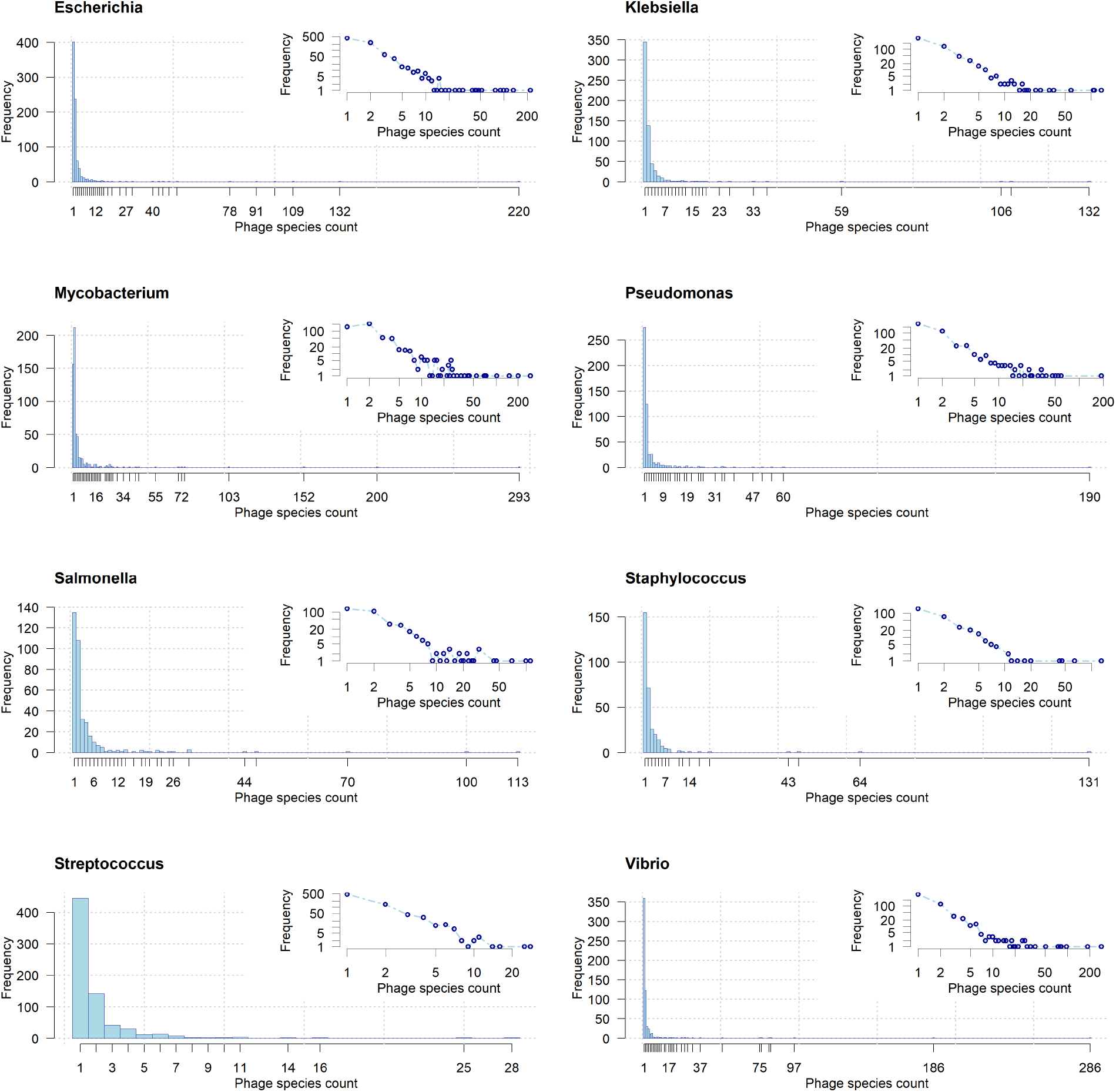
Frequency distributions of the number of times phage species have been observed by their host organisms (Escherichia, Klebsiella, Mycobacterium, Pseudomonas, Salmonella, Staphylococcus, Streptococcus, and Vibrio). Data from the 2025 dataset. Insets: same distributions on log-log axes.

While we focus on the number of species found in samples and subsamples, further measures of overall species diversity describing the underlying species abundance distributions are interesting to discuss - see Supplementary text 4 for further summaries and analysis of species diversity.

### 3.2. Internal validation

For our internal validation, we have summarised the error distributions in Figures 2 for dataset DB25 (Supplementary Figure 4 for DB24). Table 2 and Figure 3 compares median (normalized absolute) errors between the different estimators across all training set size and host genus combinations. For all ratios between training set size and the size of the set we want to predict, we see that the PYP estimator shows, in many training set size and host genus combinations, higher errors than the other estimators, although the differences are not necessarily big. For DB25, OSW shows more often lower median errors than all other estimators (Table 2, Figure 3). This picture slightly changes for DB24 where FPG outperforms OSW measurably, while CJ shows only very slightly better performance (Table 2, Figure 3). For the two overall best estimators OSW and FPG, we also considered an alternative prediction scenario for DB25, from fixed training set sizes (200, 300, …, 500) the number of additional species of a further set of samples of smaller or larger size (100, 200, …, 500). In this validation setup, OSW performs slightly, but nearly consistently better (Supplementary Text 3). Only for training set size *n* = 200 for host genus *Streptococcus* FPG performs slightly better when predicting a further sample of size *m* = 400, 500.

**Figure 2:**
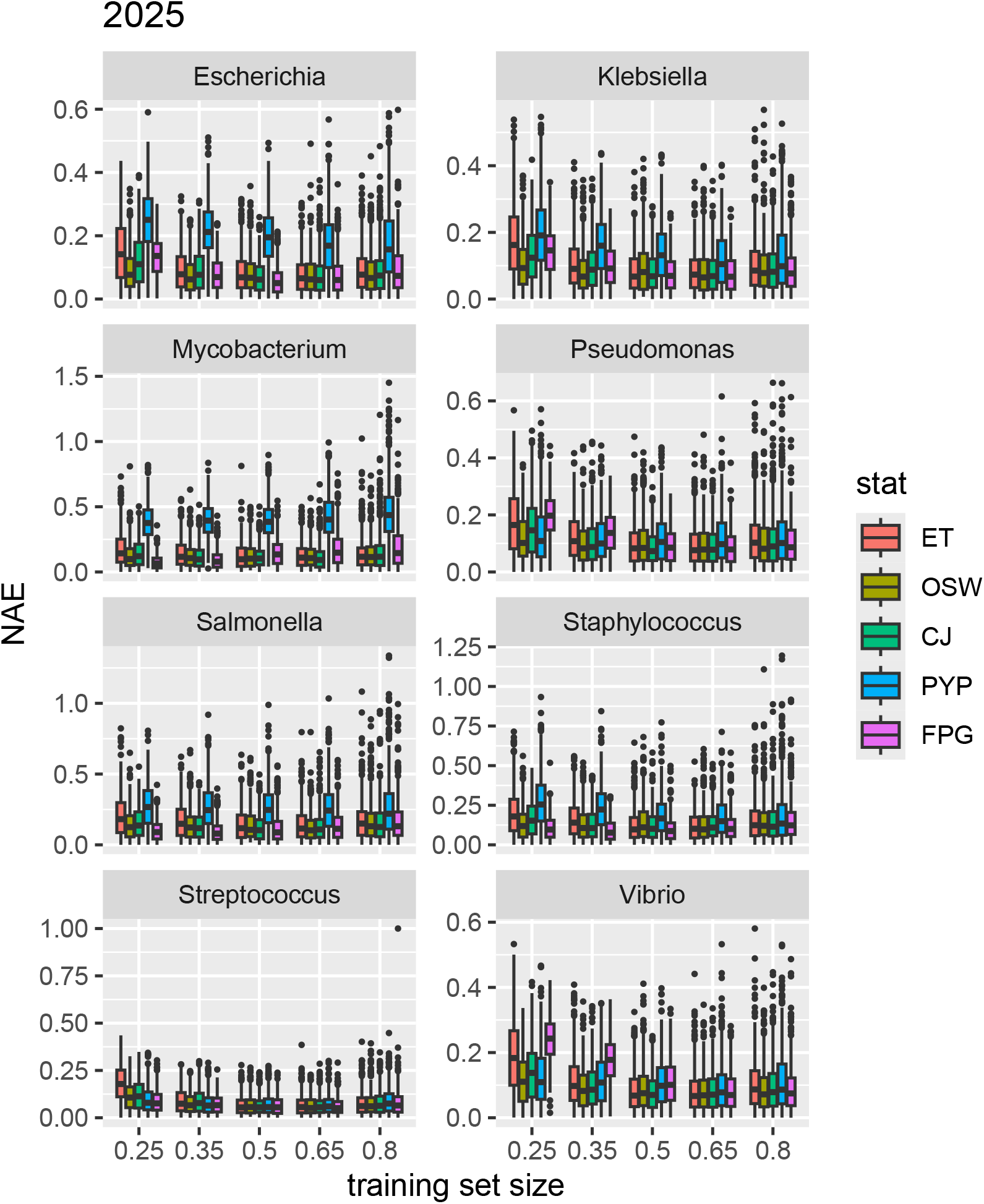
Normalized absolute errors (NAEs) for prediction of the number of unseen phage species per host genus in the rest of the database from a training set of size x% of all sampled phages from this host (DB25). Estimators CJ, FPG and OSW show comparable errors across host genera and training set/prediction set ratios.

**Table 2:** best: Counts of times each estimator had the lowest median error across training set sizes and host genera, vs OSW: Percentage of OSW having lower median error than other estimators across of training set size and species combinations when compared with each other estimator (in parentheses: percentage of comparison significant at 5%, two-sample one-sided Wilcoxon test for lower error in OSW, with Holm correction for multiple testing across all combinations). OSW, FPG and CJ perform best generally and comparably. For the comparison based on DB25, the Wilcoxon test for smaller errors for FPG than for OSW was significant 37.5%, for CJ 7.5%.

| Estim. | best: DB24 | best: DB25 | vs OSW: DB24 | vs OSW: DB25 |
| --- | --- | --- | --- | --- |
| ET | 1 | 6 | 80% (40%) | 80% (35%) |
| PYP | 0 | 0 | 97.5% (80%) | 92% (80%) |
| CJ | 10 | 5 | 47.5% (22.5%) | 62.5% (17.5 %) |
| OSW | 9 | 17 | - | - |
| FPG | 20 | 12 | 35% (20%) | 60% (27.5%) |

**Figure 3:**
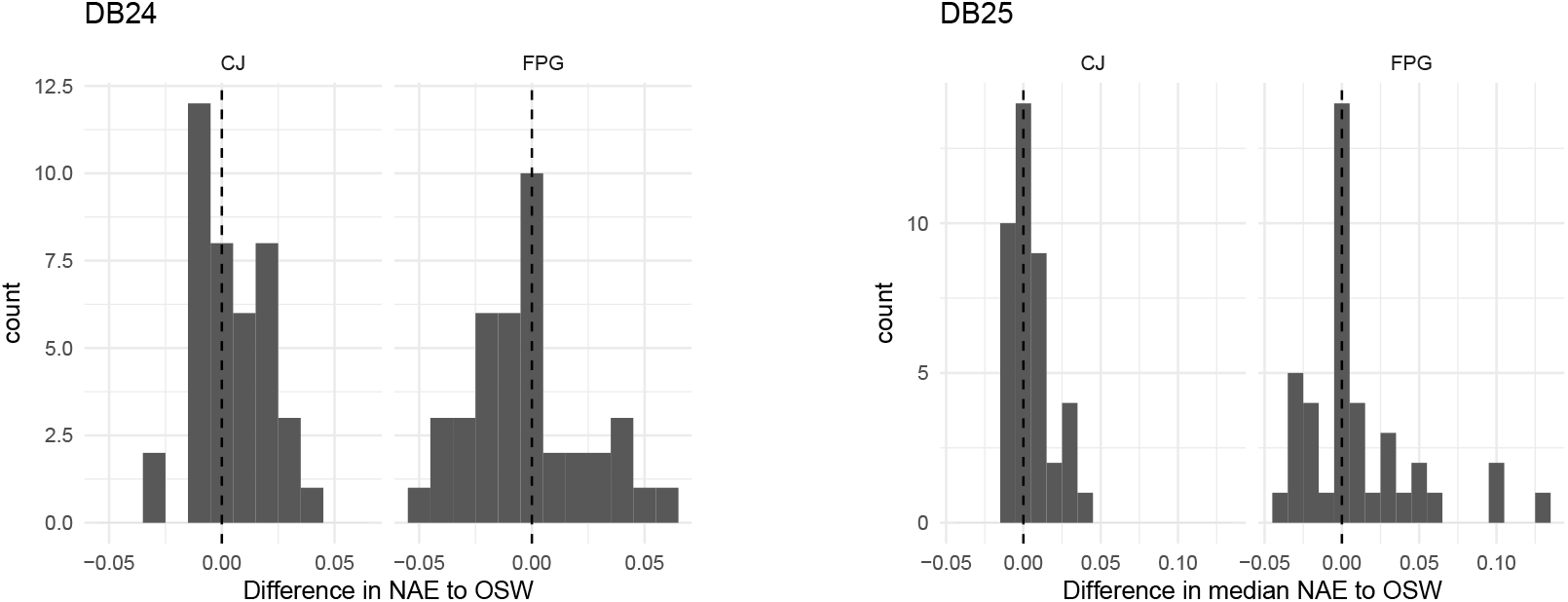
Histogram of the difference between median normalized absolute errors for CJ or FPG to those of OSW. Positive differences correspond to smaller errors of OSW.

Taken together, FPG and OSW perform similarly with CJ not fa xr behind (see also Figure 3), we would recommend OSW to predict as FPG shows considerably large errors for some combinations for DB25 (e.g., host genus *Vibrio* for low training set sizes). In many conditions, apart from PYP, all estimators perform similarly well. Thus, the choice of specific estimator among CJ, FPG and OSW is not that important. We see that, at least in our internal validation scheme, we can predict the additional species with good to medium accuracy for our best estimators. For instance, for fixed ratios between training and to-be-predicted sets, OSW shows a maximum median error across all conditions of 14.1 % for DB24 and 13.2 % for DB25 (3rd quartile: 24.8 % for DB24, 22.3% for DB25).

While errors are higher when predicting larger samples from a small training set and lower for larger training set sizes, the trends are not monotonically decreasing as illustrated in Figure 4 (DB25) and Supplementary Figure 5 (BD24).

**Figure 4:**
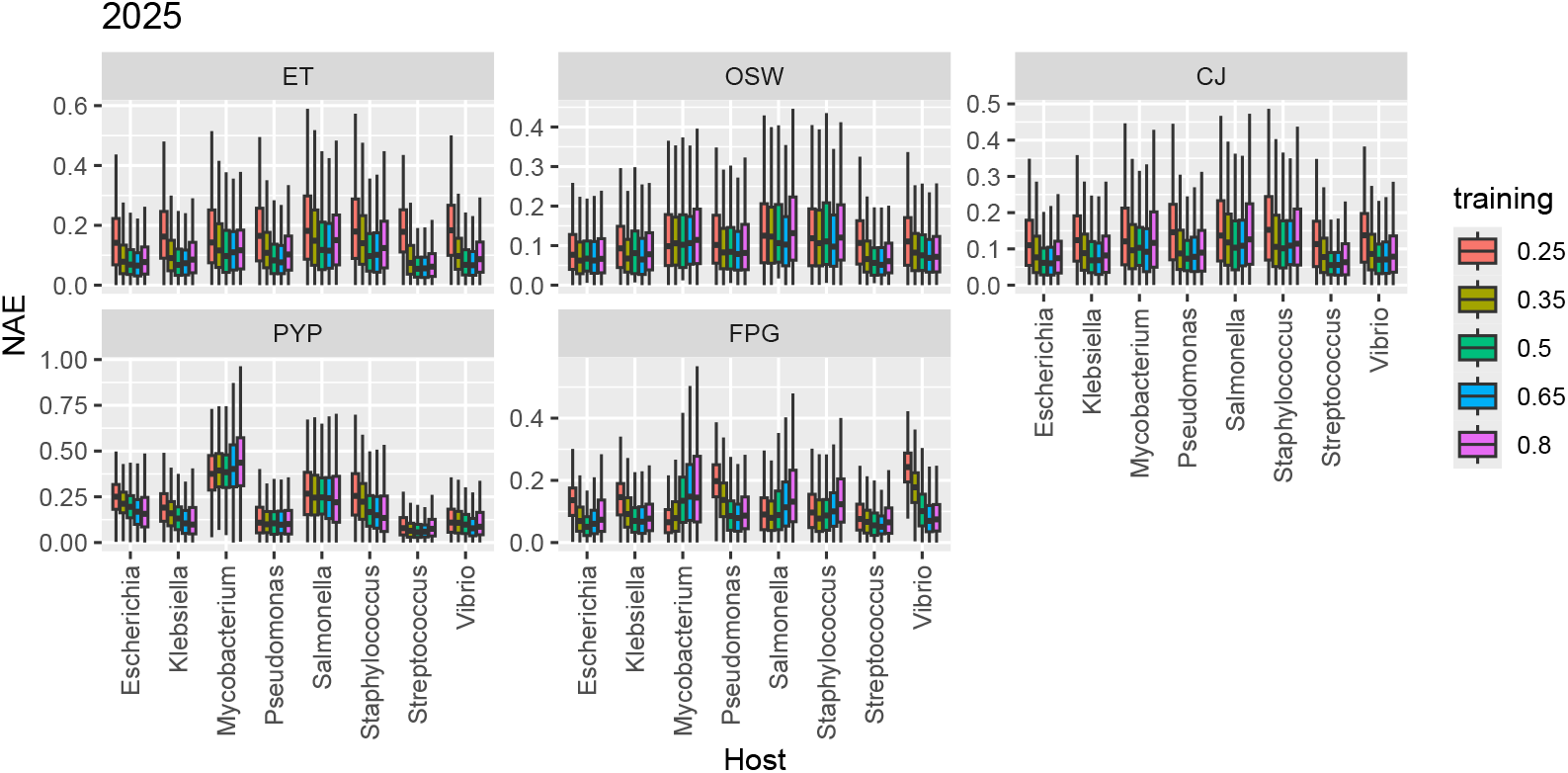
Normalized absolute errors (NAEs) for prediction of the number of unseen phage species per host species in the rest of the database from a training set of size x% of all sampled phages from this host (DB25). Outliers not shown.

### 3.3. Predicting real data - from temporal snapshot to temporal snapshot

As discussed above, all estimators are based on assuming that the species abundance distribution, or more precisely the entry distribution into the database, stays stationary over time. However, we observe that predicting the additional species in the later state of the data (DB25) from the earlier species distribution (DB24) had much higher normalised mean errors (Table 3) than predicting the species of an equally sized set from a random split of the entries in DB25 (following our internal validation approach). Moreover, apart from host *Vibrio*, estimating the later state of the database from an earlier snapshot strongly underestimated the number of newly observed phage species (negative normalised mean errors in Table 3). Comparison with the appropriate analogous subsetting for internal validation yielded that, apart from host *Mycobacterium*, these errors were also at the highest quantiles of the error distribution when comparing predicting from a random subset of equal sizes (Table 4), for all estimators but PYP. This hints at the possibility that the data generation changed between the time points. As PYP estimates were much noisier than the other estimates, the less clear picture for PYP will be, at least partially, due to the higher prediction errors.

**Table 3:** Normalised mean errors (*predicted*−*observed*)*/observed* of predicting the number of additional phages in a later database snapshot (DB25) from the species distribution at an earlier timepoint (DB24). For all host genera but Vibrio, we see underprediction of the number of additional phage species.

|  | OSW | ET | CJ | FPG | PYP |
| --- | --- | --- | --- | --- | --- |
| *Escherichia* | -0.69 | -0.69 | -0.69 | -0.67 | -0.60 |
| *Klebsiella* | -0.32 | -0.32 | -0.32 | -0.31 | -0.23 |
| *Mycobacterium* | -0.67 | -0.67 | -0.67 | -0.66 | -0.48 |
| *Pseudomonas* | -0.58 | -0.58 | -0.58 | -0.57 | -0.51 |
| *Salmonella* | -0.61 | -0.61 | -0.60 | -0.57 | -0.45 |
| *Staphylococcus* | -0.61 | -0.61 | -0.61 | -0.60 | -0.55 |
| *Streptococcus* | -0.66 | -0.66 | -0.66 | -0.64 | -0.62 |
| *Vibrio* | 0.84 | 0.74 | 0.88 | 1.02 | 1.42 |

**Table 4:** Quantile of normalized absolute error when predicting the additional species in a database snapshot (DB25) from the species distribution at an earlier timepoint (DB24) in the null distribution of the comparable random set prediction described above. For all host genera but *Mycobacterium*, we see a tendency to see uncharacteristically high errors for the prediction of DB25 from DB24, indicating changes in the underlying species abundance distributions entering the database.

|  | OSW | ET | CJ | FPG | PYP |
| --- | --- | --- | --- | --- | --- |
| *Escherichia* | 1.00 | 1.00 | 1.00 | 1.00 | 0.98 |
| *Klebsiella* | 0.98 | 0.98 | 0.98 | 0.97 | 0.83 |
| *Mycobacterium* | 0.70 | 0.74 | 0.69 | 0.73 | 0.51 |
| *Pseudomonas* | 0.99 | 0.99 | 0.99 | 0.99 | 0.98 |
| *Salmonella* | 0.96 | 0.96 | 0.97 | 0.94 | 0.75 |
| *Staphylococcus* | 0.95 | 0.94 | 0.94 | 0.96 | 0.87 |
| *Streptococcus* | 1.00 | 1.00 | 1.00 | 1.00 | 1.00 |
| *Vibrio* | 1.00 | 1.00 | 1.00 | 1.00 | 1.00 |

This analysis shows that, for all host genera but *Mycobacterium*, the underlying database entry distribution, and thus very likely the phage sampling strategies, have not been stable over time.

### 3.4. Predicting the efficiency of future phage sampling

To predict the number of new phage species to be discovered in future samples, the presented approaches need stationarity of the underlying species distribution of database entries, i.e. that the species entering the database come from a species distribution not changing over time. If this assumption holds, indeed, we can use the presented estimators and would expect manageable prediction errors, based on our validation results. However, our temporal analysis indicates that only for host genus *Mycobacterium* we can assume this has not changed over time. This means that prediction for the other host genera is only valid if future samples come from the now updated species distribution that mixes different distributions from different time-points - an assumption that does not necessarily hold.

Under this stationarity assumption, what do the estimators predict for future sampling efforts? Figure 5 shows the predictions for (the monotone-concave modification of) OSW, FPG and CJ. Predictions differ between estimators, but qualitatively agree for host genera *Klebsiella, Pseudomonas, Staphylococcus, Streptococcus* and *Vibrio*. For these host genera, if the entries into the database follow the distribution of the previous ones, we can expect a linear gain for small additional sample sizes and then a slightly sublinear gain if *m* increases. When projecting the number of sampled individuals to twice the current number of isolates, we predict at least 229 additional species (≈ 37 % of currently observed species) for *Klebsiella*, 158 (31%) for *Pseudomonas*, 96 (31%) for *Staphylococcus*, 321 (46%) for *Streptococcus*, and 247 (41%) for *Vibrio*. For host genera *Escherichia, Mycobacterium* and *Salmonella*, the prediction of estimators diverge: OSW predicts that *de facto* saturation may be reached for all three species, CJ only for *Mycobacterium* and *Salmonella* (on slightly higher level), while FPG predicts that there will still be a steady discovery of new species in new samples. These indicated saturations/diminishing returns would slow species gain so strongly that they even fall below extrapolating a *#*species ∼ *log(*sample size) model fitted to the species accumulation curve (following [23], see Supplementary text 2 and Supplementary Figure 8 within) for *Mycobacterium* and *Salmonella*, which provided a clear lower bound for all estimators for other genera (and for the true species in internal validation when predicting from 80% of all samples). However, these estimates are only reliable if sampling efforts and data base entry strategies stay comparably to past efforts - which as we saw is not a given for phage databases.

**Figure 5:**
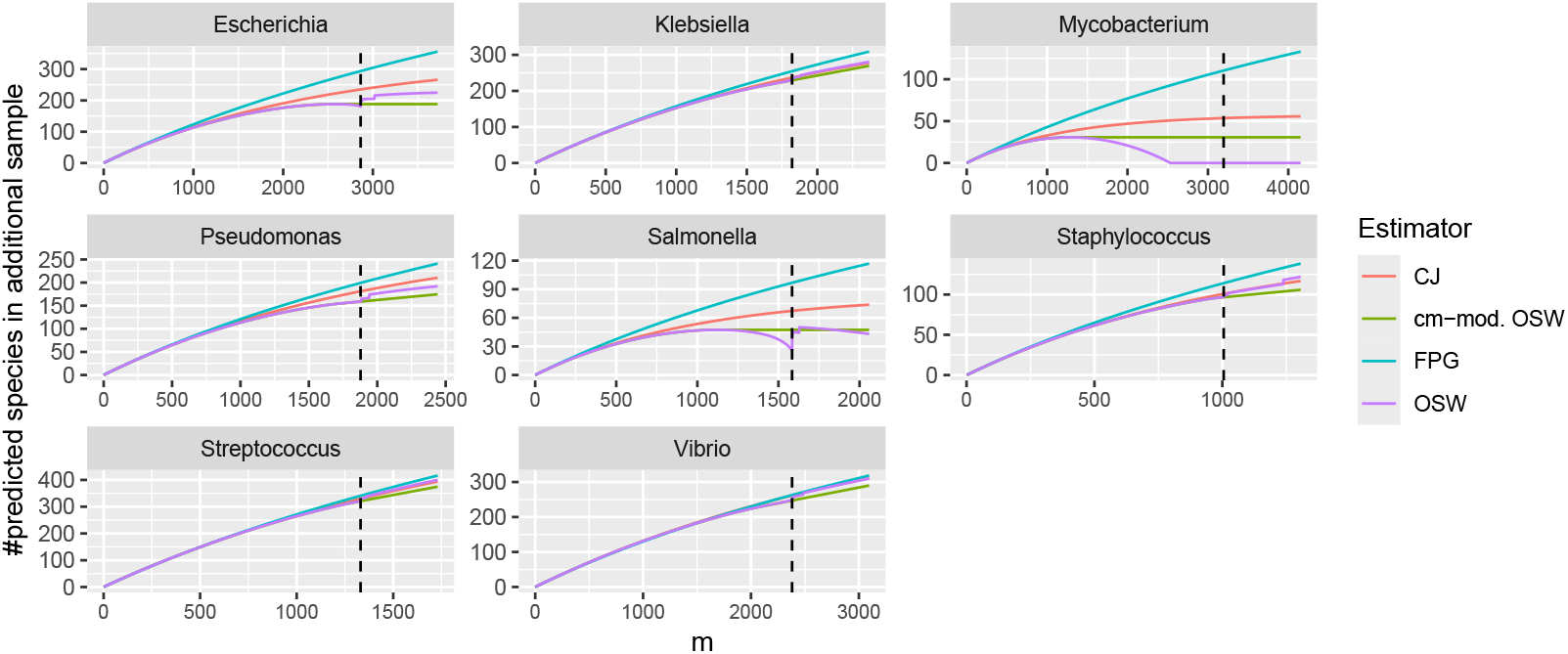
Prediction of the number of additionally observed species by OSW (with and without concave-monotonic modification), CJ and FPG in an additional phage sample from the host genus of size *m*. Dashed vertical line show the current number of phages in the DB25 snapshot. Estimations are based on the DB25 snapshot. See Supplementary Figure 6 for added bootstrap quantiles.

## 4. Discussion

To provide sustainable control of bacterial pathogens by phages one needs to be able to estimate diversity of phages from limited samples. Here we explore how well we can predict the number of new phage species to be potentially discovered from new samples, when *m* more individual phages are isolated. In this study, we assessed five established SSP estimators for the number of unseen species. We focused on the phage species distribution for eight common bacterial genera from a well-curated phage genome database, and evaluated the accuracy of these estimation procedures using a subsampling approach to extract training and validation sets from the database. As we do not have any mechanistic theories about phage speciation and species distributions, such validation approaches, i.e. approximating the unknown species abundance distribution through subsampling is essentially the only option to assess the distribution from the current state of sampling. However, such approaches, are naturally dependant on whether the already collected samples are representative of the underlying species distribution [21]. On top, any species sampling question also depends on what counts as a species. i.e. how we delineate species. We use a definition for phage species based on sequence similarity and coverage, with a typical similarity threshold as adopted in the phage genomics community [24, 25].

Our analysis sheds light on the difficulties of predicting the efficiency of future species sampling, but allow some pointers towards analysing species sampling strategies. First, we show that several estimators predict similarly well (in our internal validation scheme), with only the parametric Pitman-Yor-prior-based PYP estimator, the computationally most expensive estimator, trailing clearly behind the other estimators. This may be due to a sub-optimal fit of the Pitman-Yor distribution to the underlying species distribution *per se*, but also to known issues with large-sample consistency as reported in [12]. We have not explicitly discussed methods based on simply extrapolating the rarefied species accumulation curve, as a preliminary test of one such approach (which approximates this curve by a semilog model [23]) showed, for our data, comparably higher errors than the methods presented here (Supplementary text 2).

Using the non-parametric OSW or CJ predictors or the parametric FPG estimator for the number of unknown species leads to controllable errors when predicting the number of additional species (for OSW, errors typically, i.e. their third quartile, stay below 25% of the current value). However, these error estimates are only directly indicative of estimation precision of future samples if the species abundance distribution of the phages entering the database does not change over time. This is not only critical for our internal validation scheme, but a critical assumption behind all estimators of additional species found in future samples we used and discussed here. Their statistical model assumes a common underlying species abundance distribution, i.e. its stationarity for all samples, present or future [10, 22, 26]. However, for all host genera but *Mycobacterium*, our analysis of predicting a later sampled state (DB25) from a previous state (DB24), reveals that sampling, or more precisely database entry species abundances have changed over time, questioning this crucial assumption. Indeed, for all genera but *Mycobacterium* and *Vibrio*, all estimators underpredicted the actual new species entering the database, which is often reported for estimators of additional species in future samples [23, 21]. Interestingly, host genus *Vibrio* shows an overprediction. In terms of sampling and database entry efforts for phages, there is an interpretation of the differences between genera: Most genera have been more variably sampled over time, with sampling from an increasing variety of host species and environmental contexts over time. A notable exception is *Mycobacterium*, whose phages have been sampled predominantly from host species *Mycobacterium smegmatis* as part of the SEA-PHAGES phage isolation program [27]). This strategy for sampling has not changed during collection time reflected in the database. This may also explain the potential saturation after additional sampling (Figure 5), as the range of host species for *Mycobacterium* is much narrower than for other bacterial host genera considered. For *Vibrio*, on the other hand, phage sampling (and subsequent database entry) reportedly shifted to a previously less sampled host group (phylogenetic clade) [25] - thus highlighting a potentially new but narrow set of hosts, leading to a more narrow/less rich species abundance distribution - which may explain the observed overprediction.

Taken together, our findings suggest that there is a difference between the underlying species distributions of samples entering the database at different time points. This has a clear implication for how predictions from the database on future sampling should be interpreted: we predict how many new phage species will enter the database under the assumption that new isolates are drawn from the same underlying species distribution as the current data in other words, if the sampling and database entry strategy does not change when sampling new isolates. As it has likely changed previously, this does mean that predicting the number of additional phages in future samples using any of our estimators is only reliable if the new sampling strategy is a similar mix of strategies as used until now.

Under this constraint, we still may cautiously interpret our prediction results. We would expect, if future sampling strategies are mixed similarly between the sampling efforts behind previous database entries, that for host genera *Klebsiella, Pseudomonas, Staphylococcus, Streptococcus* and *Vibrio*, future sampling will provide ample new species. There is some evidence for host genera *Mycobacterium* (where predictions are more reliable as we have no statistical evidence that sampling strategies have changed), *Escherichia* and *Salmonella* that new phage species found may plateau with future sampling efforts. For the latter three species we found that predictions are also estimator-dependent, as also reported for other data sets [21]. We emphasise that predicting from the database can and will only predict how many phage species enter the database, as this is the distribution we have information on - which reflects the true underlying ecological species abundance distribution only through the lens of research and curation bias. Differences in the rate at which we may discover additional species in future samples for different host genera are expected. The genera differ in the mixture of ecological niches for their member species, their biological and genetical properties, which may affect the inclusion and selection for or against phages, but also different sampling strategies.

For further assessment of the differences in species abundance diversity within the database, as we here were only directly interested in the number of different species, see Supplementary text 4.

Our analysis focussed on the eight most common host genera, and can be easily applied to other host genera, given enough sampled phages. However, can we also predict the number of phage species for an unsampled host or a metagenomic environmental sample with multiple different hosts? Our strategy this would be to generate an expected distribution for this by simply sampling the appropriate size from a random genus in the database (in the case of a single host genus/species) or generally sample appropriately from the full database for a given number of random hosts predicted to be present within an environmental sample.

A different limitation arises from the aggregation of phages across host genera: phages infecting hosts of the same genus will not be infecting all bacterial species within that genus - thus, for a direct application our prediction may be misleading as we have not predicted which fraction of phage species collected from a host genus affects a given species. However, if we can increase the resolution to host species or estimate this fraction, or alternatively consider regional or sampletype sampling sets, the estimation approach would remain also as valid at these levels - it could prove even more reliable, as on these levels, sampling strategies may have been more stable. However, at least directly from the database, we do not have these resolution for our samples. Still, we still can predict the efficiency of future sampling, just not whether all these species would be useful to combat specific bacterial species.

In conclusion, our study highlights a general need for adjusting phage sampling strategies in order to maintain a steady inflow of new phage species. Our analysis shows that for host genus *Mycobacterium*, future sampling efforts with the same sampling strategy may prove inefficient, while for all other genera observed we could detect that there is evidence for a paste change in sampling respectively data base entry strategy, which lead to increasing species diversity in genera *Klebsiella, Pseudomonas, Staphylococcus, Streptococcus, Salmonella* and *Escherichia* over time, but a more narrow species range than earlier for *Vibrio*. Due to that change in sampling strategies, prediction methods of additional species in further phage samples entering the data base, developed for the classic species sampling problem, have limited prediction ability and their results should be interpreted with caution.

## 5. Data and code availability

All data is available via the INPHARED database https://github.con/RyanCook94/inphared, https://nillardlab.org/bacteriophage-genonics/inphared/. The analysis scripts have been made available at https://github.con/ncavallaro/PhageDiversity.

## 6. Acknowledgements

This project was supported by a seed corn grant of the Leicester Institute for Advanced Studies (LIAS), and by our colleagues: Christopher Bayliss, Sarah Diver, Jason Hughes, Eva Krockow, Sergei Petrovskii, Andy Schofield, Carolyn Tarrant from the interdisciplinary phage research group of the University of Leicester. AK was supported by the Medical Research Council [grant number MR/W007002/1]. Andrew Millard was supported by the Medical Research Council MR/T030062/1 and MR/L015080/1, United Kingdom. Andrew Morozov supported by the Engineering and Physical Sciences Research Council (EPSRC, EP/W522326/1) of the United Kingdom. We thank three anonymous reviewers for their constructive feedback.

