## Supplementary texts and figures for "How many phage species remain undiscovered? Species sampling approaches to inform phage discovery"

#### Contents

**F1: Supplementary Figures** **2**

**1 Bootstrap approach for the ET estimate** **8**

**2 Baseline comparison for the number of species in a sample via a semi-log linear model** **10**

**3 Internal validation from fixed sample sizes to fixed new sample size** **11**

**4 Exploring phage diversity using Hill numbers** **29**

4.1 Diversity metrics - methods . . . . . . . . . . . . . . . . . . . . . . 29

4.2 Phage diversity - results . . . . . . . . . . . . . . . . . . . . . . . . 29

### F1: Supplementary Figures

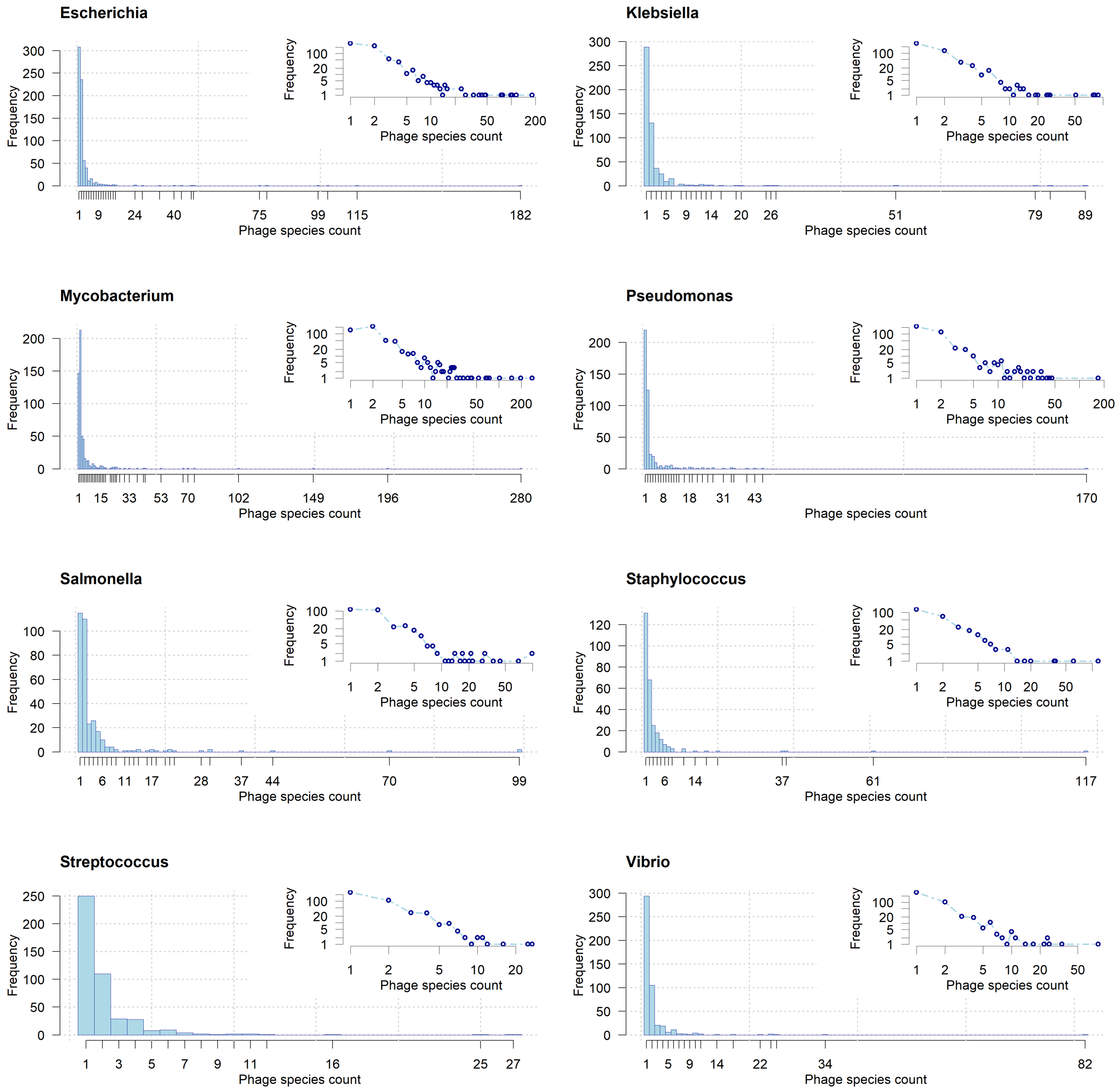


Figure 1: Frequency distributions of the number of times phage species have been observed by their host organisms (*Escherichia*, *Klebsiella*, *Mycobacterium*, *Pseudomonas*, *Salmonella*, *Staphylococcus*, *Streptococcus*, and *Vibrio*). Data from the 2024 dataset. Insets: same distributions on log-log axes.

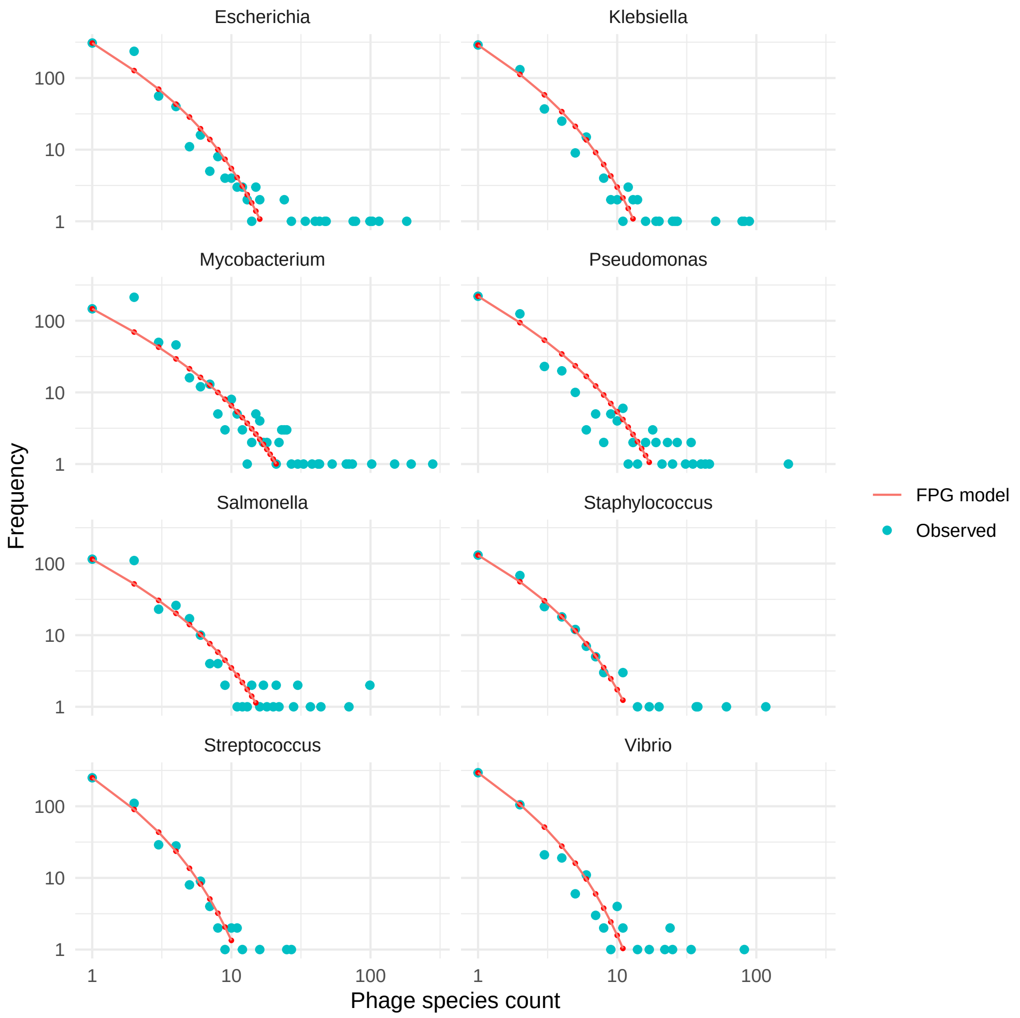


Figure 2: Comparison of observed species abundance counts with expected species abundance counts under a mixed Poisson-Gamma distribution (drawing as many individuals as phages sampled) with best (ML) fitting parameters to the observed values

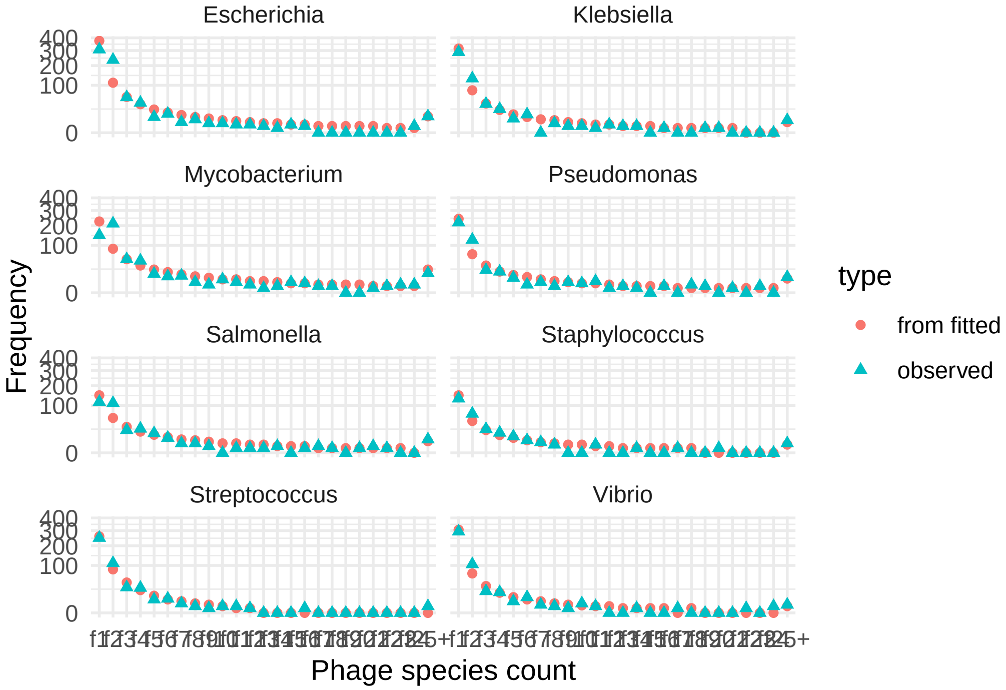


Figure 3: omparison of observed species abundance counts with median species abundance counts simulated 500x under a Pitman-Yor distribution (drawing as many individuals as phages sampled) with best (ML) fitting parameters to the observed values

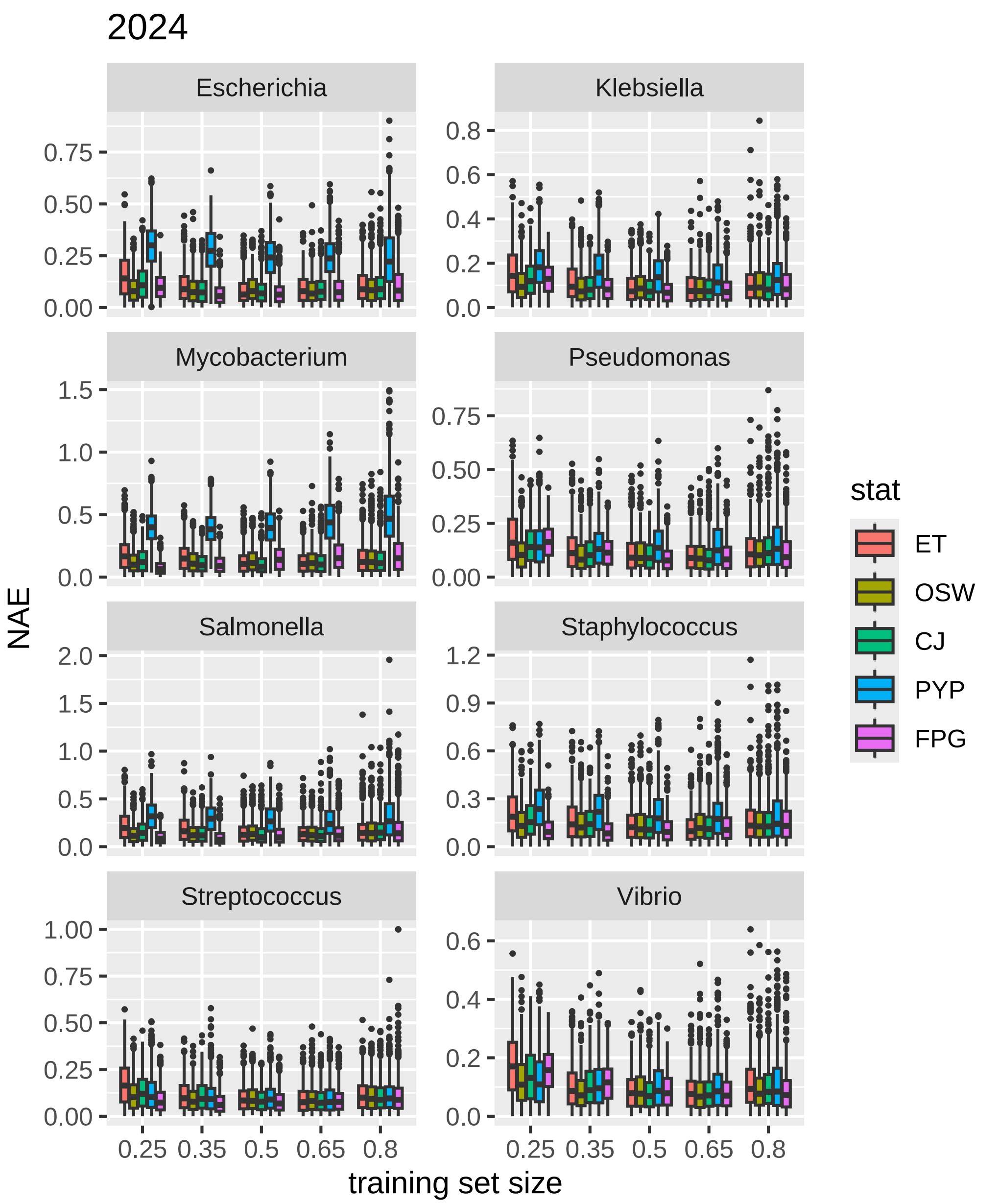


Figure 4: Normalized absolute errors for prediction of the number of unseen phage species per host genus in the rest of the database from a training set of size x% of all sampled phages from this host (data basis: snapshot from September 2024, DB24)

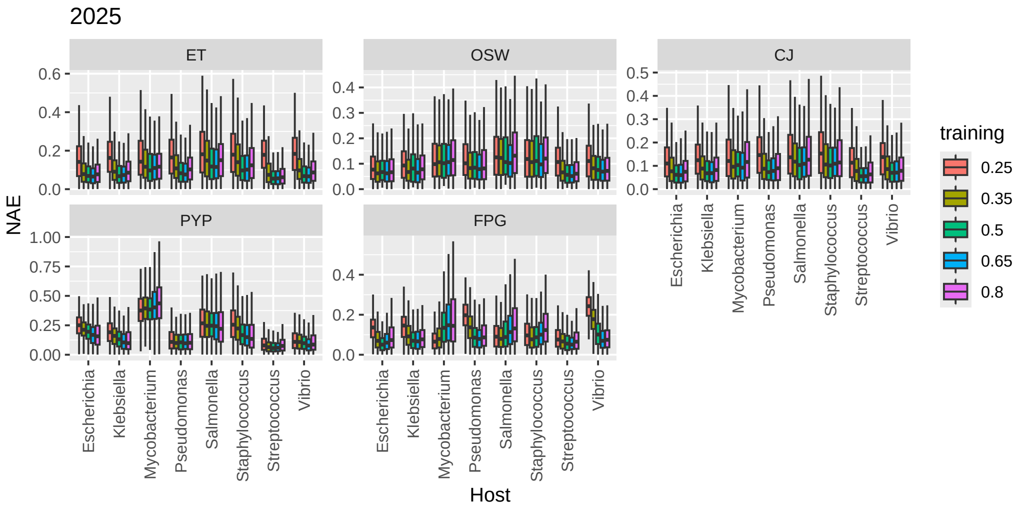


Figure 5: Normalized absolute errors for prediction of the number of unseen phage species per host species in the rest of the database from a training set of size x% of all sampled phages from this host (data basis: snapshot from September 2024, DB24). Outliers not shown.

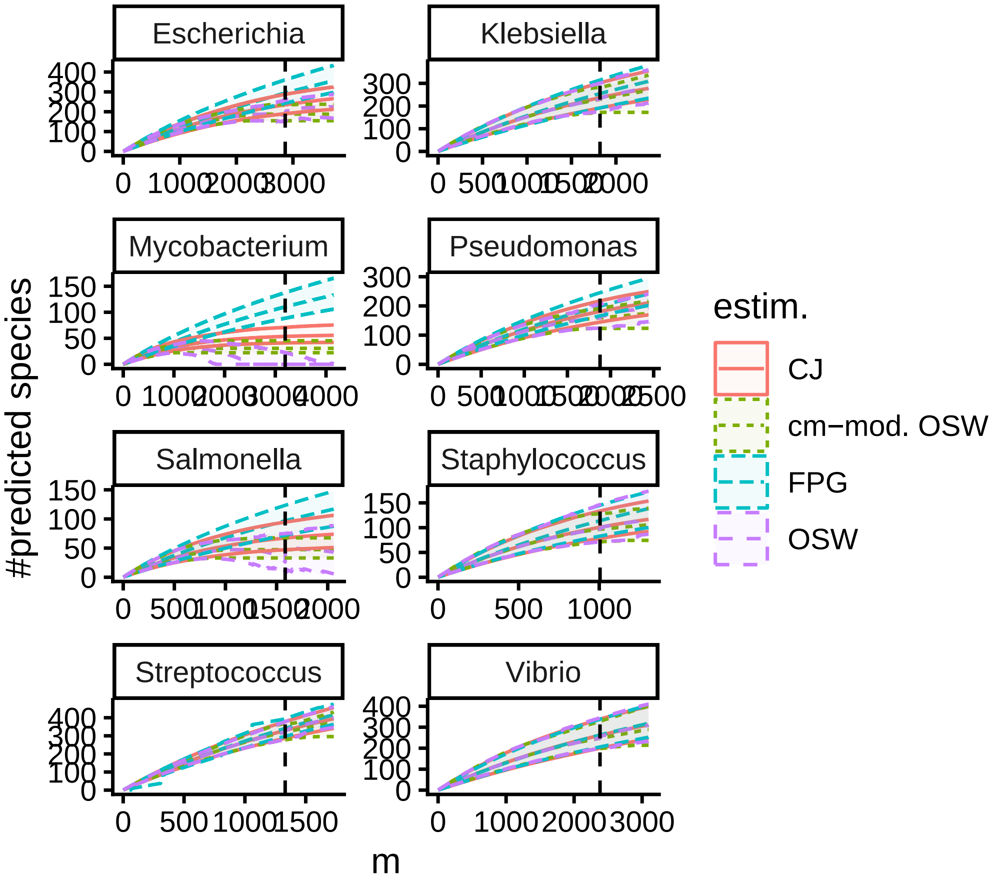


Figure 6: Prediction of the number of additionally observed species by OSW (with and without concave-monotonic modification), CJ and FPG in an additional phage sample from the host genus of size $m$, with surrounding bootstrap 2.5% and 97.5% quantiles (100 bootstrap replicates). Dashed vertical line show the current number of phages in the DB25 snapshot. Estimations are based on the DB25 snapshot.

### 1 Bootstrap approach for the ET estimate

Figures 7 show species-based and individual-based bootstrapping (10 bootstrap samples each) for three random subsets of phage sets for the genus *Escherichia* from snapshot DB25. The distribution of ET estimates from bootstrap samples of a specific subset of size 250 closely resembles the distribution of ET from random subsets of the same size, which justifies this approach. We refrained from bootstrapping on the individual phage level, as such a bootstrap sample usually loses species and cannot represent real species distribution.

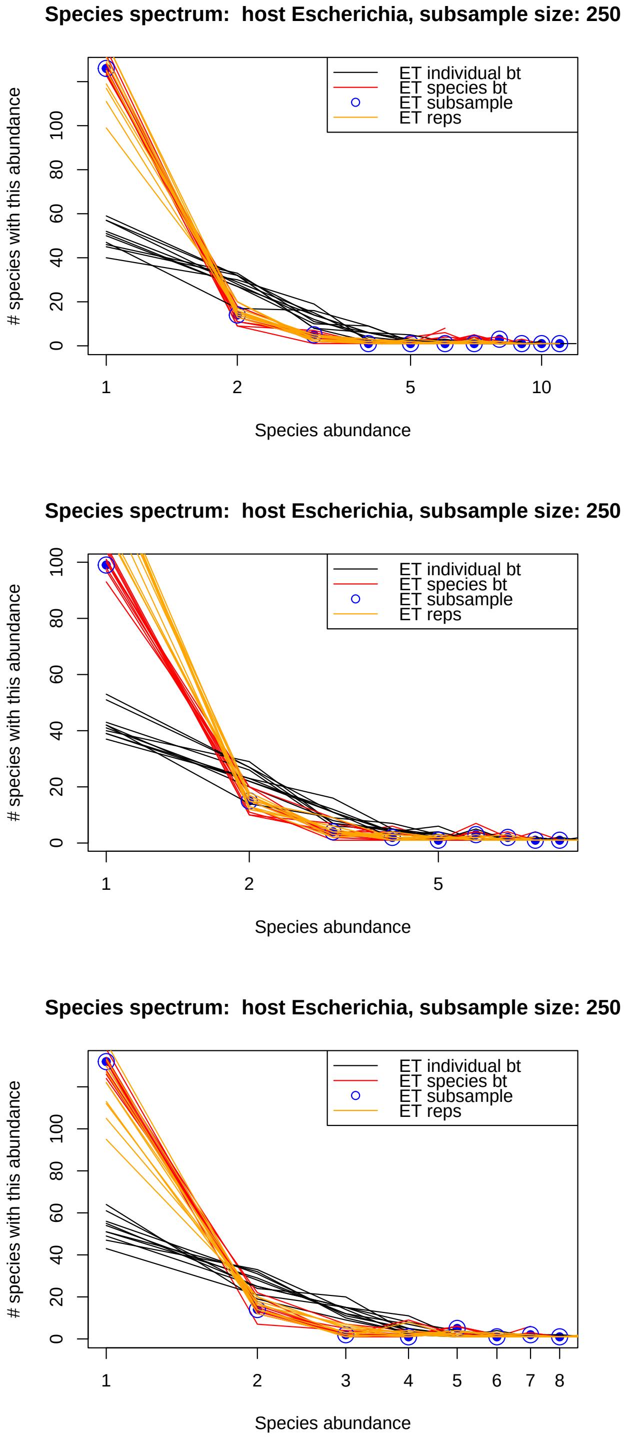




Figure 7: Comparison between species frequencies obtained from species-based and individual-based bootstrap sampling.

### 2 Baseline comparison for the number of species in a sample via a semi-log linear model

As the species sampling problem, including predicting the number of additional species found in new samples, is a well-researched statistical problem, many further estimators have been proposed, including “simply” extrapolating the species accumulation curve, i.e. how many samples can be/are expected to be found in subsamples of the dataset at hand. As a baseline comparison, we have added such an approach. We follow [1, Appendix 2], who show that for their data the first 20 species frequency spectrum entries (species that appear once, twice,...) agree well with a log-log linear model (which also holds for our genera) and then use this approximate linear model to approximate the (semi-log linear) model $\#species \sim \log(sample\ size)$ by replacing the species frequency spectrum by its log-log linear model. We use a slightly adjusted approach. As our species frequency spectra are sparse and have already counts of zeroes for species that appear exactly $i$ times in the sample for $i \leq 20$, we only fit a log-log linear model for all entries of the spectrum until the first entry with value 0.

When we assessed this approach in internal validation (predicting the missing 20% from a random subsample of size 80%), it constantly underestimated the number of species in further samples in the internal validation, including for the sample itself (additional sample size $m = 0$). Thus, we adjusted the semi-log model by adding a constant to it so that it predicts the observed number of species in the sample it is fitted to. When rerunning the internal validation, this lead to much smaller error, but still underestimation across all validation replications for all genera but *Mycobacterium*, where 98% of the internal validation runs of this semilog model with correction to match the observed species for $m = 0$ underestimates the species in the 20% remaining database entries. However, while this correction improved errors, these were much higher than errors for our other estimators: *Escherichia* 0.39-0.66 (median: 0.56), *Klebsiella* 0.47-0.69 (median: 0.61), *Mycobacterium* 0.00-0.52 (median: 0.3), *Pseudomonas* 0.35-0.72 (median: 0.59), *Salmonella* 0.06-0.65 (median: 0.42), *Staphylococcus* 0.26-0.7 (median: 0.56), *Streptococcus* 0.55-0.72 (median: 0.65), *Vibrio* 0.48-0.73 (median: 0.63). We thus refrained from further assessing this method, as for our datasets it does not show the desired level of performance, opposed to e.g. for the data from [1].

Nevertheless, as species accumulation curve extrapolation is often success-

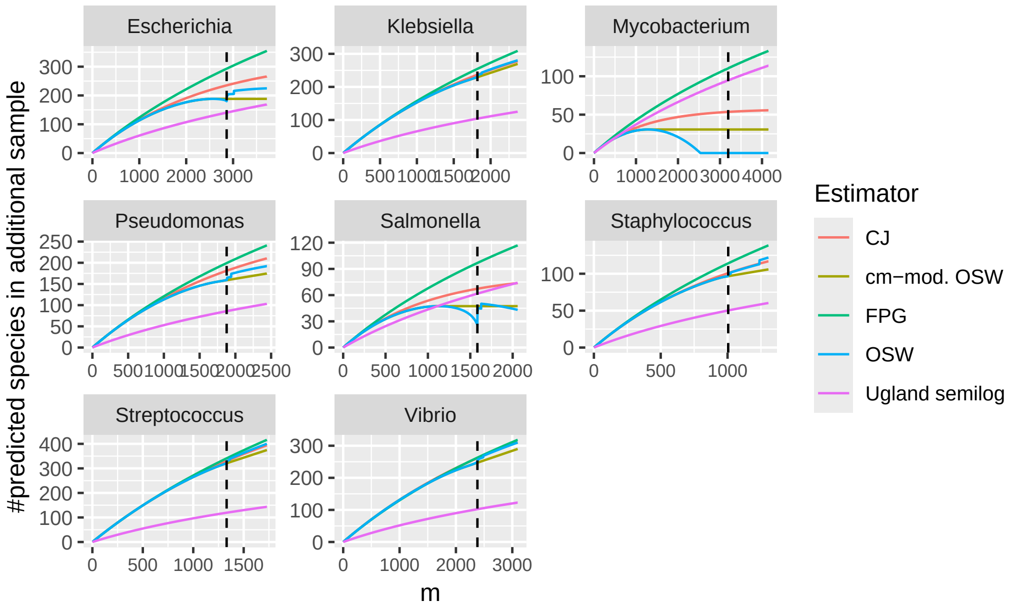


Figure 8: Prediction of the number of additionally observed species by OSW (with and without concave-monotonic modification), CJ, FPG and Ugland et al.s semi-log species accumulation curve extrapolation method, in an additional phage sample from the host genus of size $m$. Dashed vertical line show the current number of phages in the DB25 snapshot. Estimations are based on the DB25 snapshot.

fully used to predict the number of additional species in further samples, we still provide the prediction of this semi-log model for comparison to the other estimators for predicting further sampling of species from DB25 below (Figure 8).

### 3 Internal validation from fixed sample sizes to fixed new sample size

We show for each host genus considered, the normalized absolute errors for predicting the additional phages observed in a new sample of size $m$ based on the phage species distribution of already sampled $n$ phages, using the estimators OSW and FPG. First, we show all data points including outliers (outside of the boxplot whiskers, which span 1.5 IQR), then we replot without outliers.

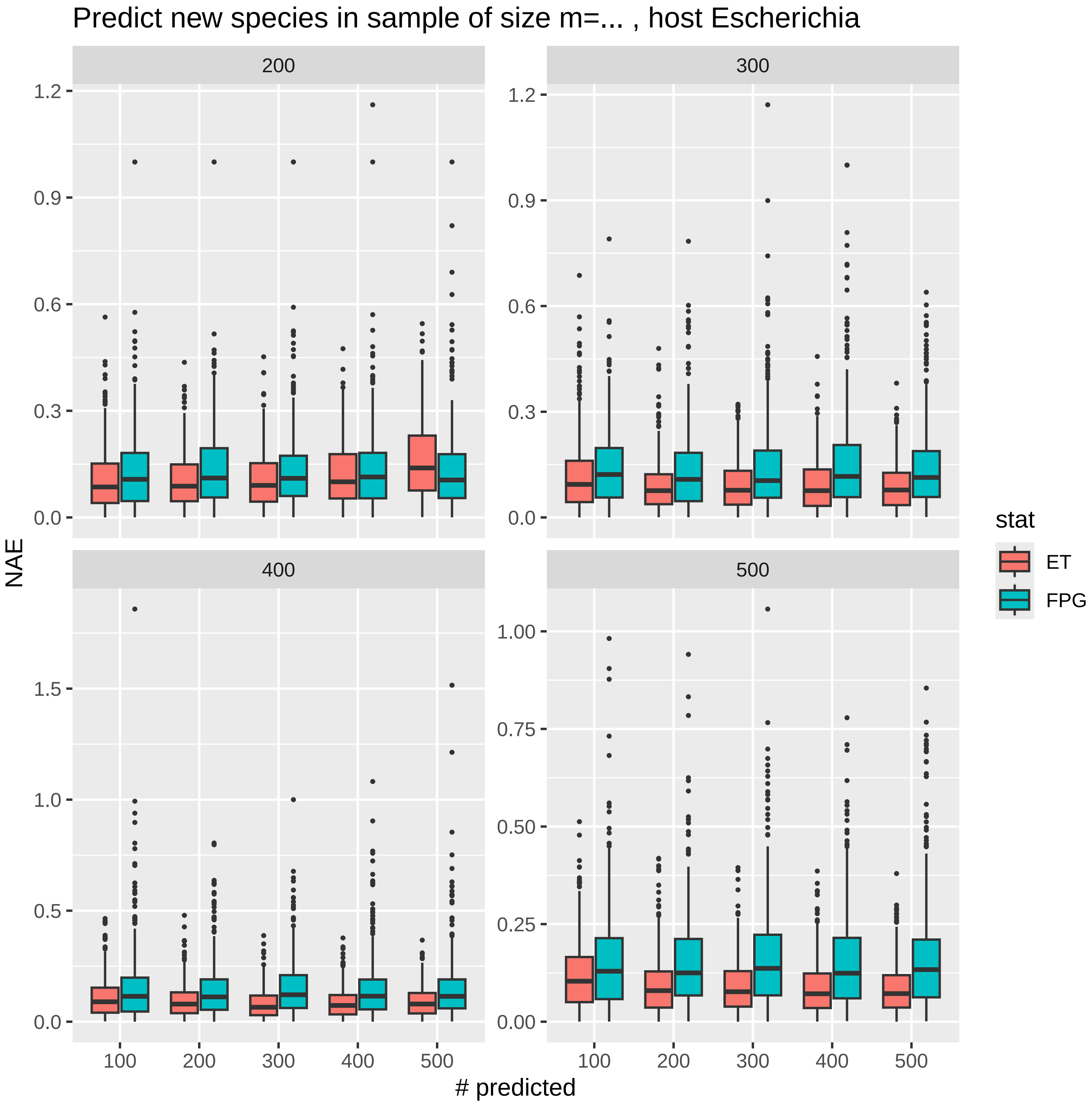

Predict new species in sample of size m=... , host Escherichia
200
300
400
500
NAE
### predicted
stat
ET
FPG
1.2
0.9
0.6
0.3
0.0
1.5
1.0
0.5
1.00
0.75
0.50
0.25
0.00
100
200
300
400
500

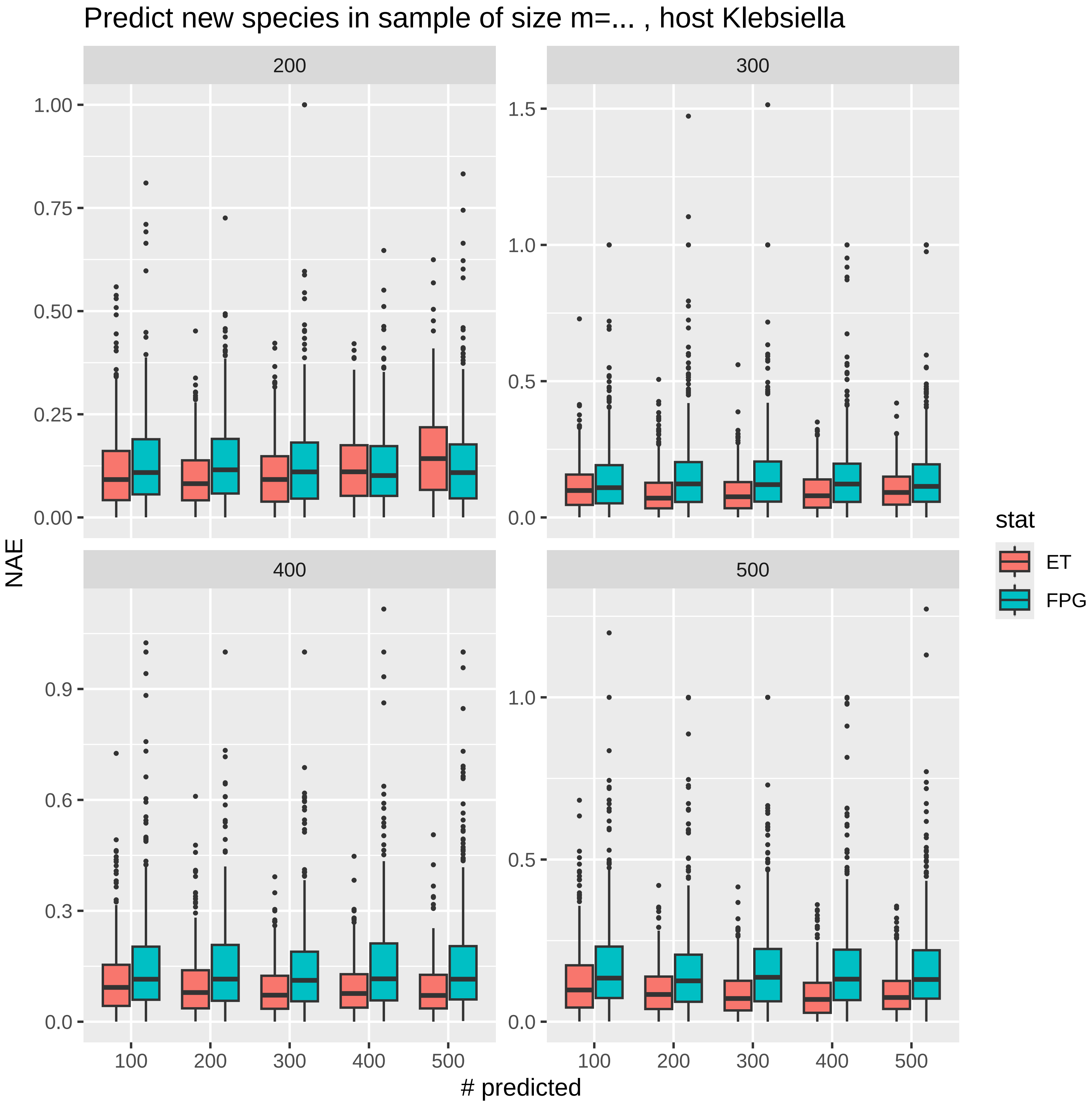

Predict new species in sample of size m=... , host Klebsiella
200
300
400
500
1.00
0.75
0.50
0.25
0.00
1.5
1.0
0.5
0.0
0.9
0.6
0.3
NAE
100
200
300
400
500
### predicted
stat
ET
FPG

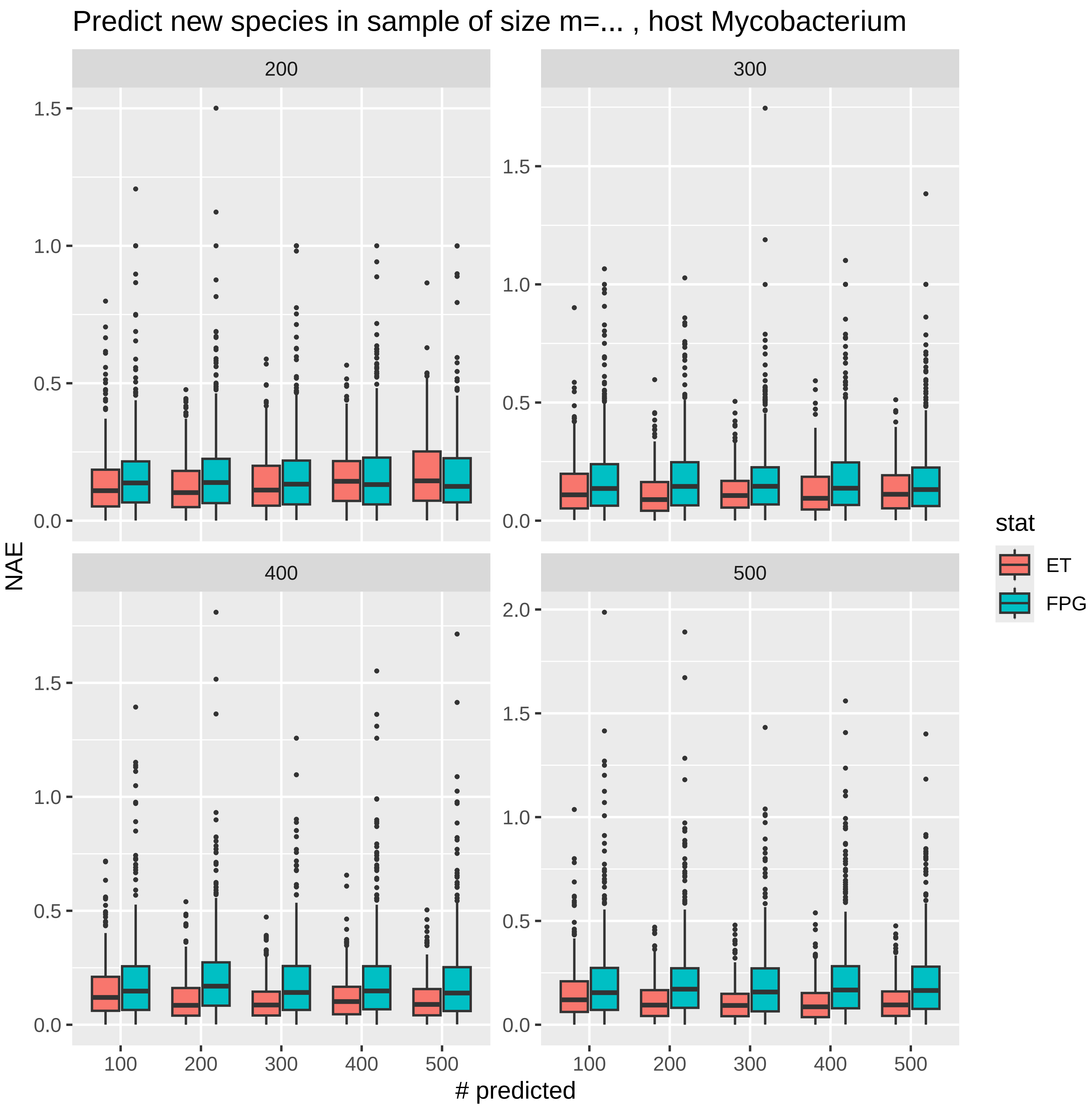

Predict new species in sample of size m=... , host Mycobacterium
200
300
400
500
1.5
1.0
0.5
0.0
2.0
NAE
100
200
300
400
500
### predicted
stat
ET
FPG

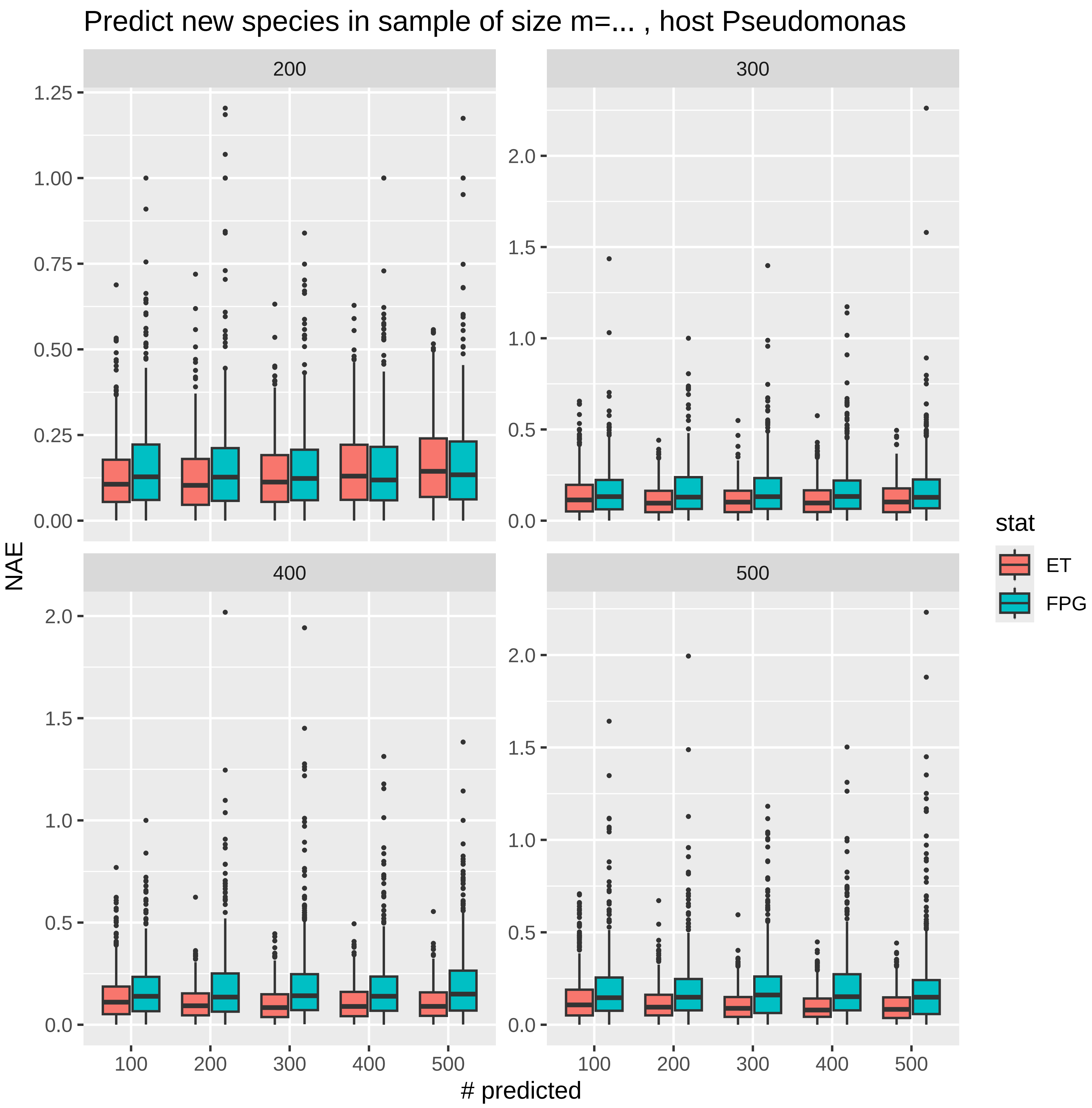

Predict new species in sample of size m=... , host Pseudomonas
200
300
400
500
NAE
### predicted
stat
ET
FPG

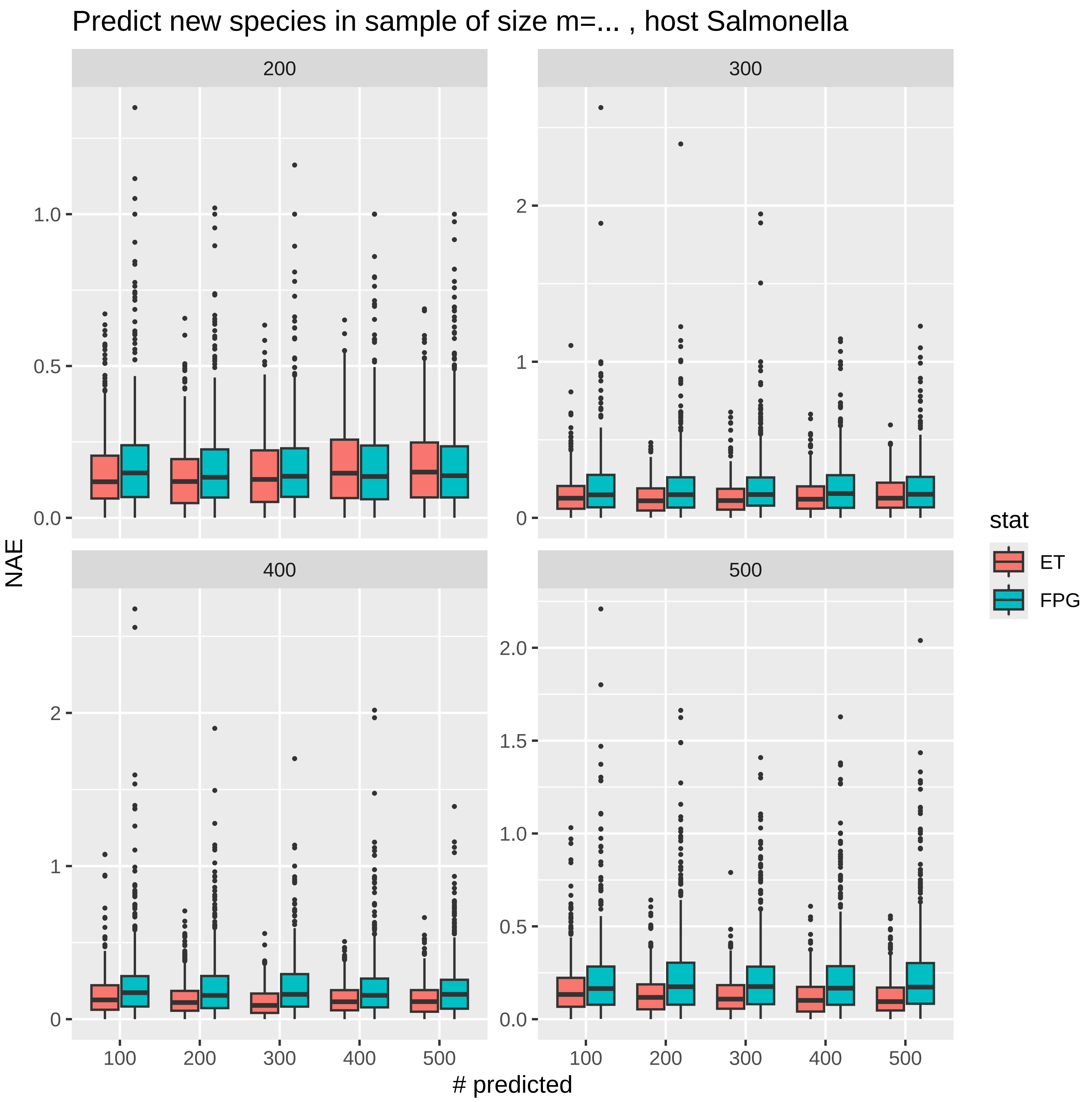

Predict new species in sample of size m=... , host Salmonella
200
300
400
500
NAE
### predicted
stat
ET
FPG

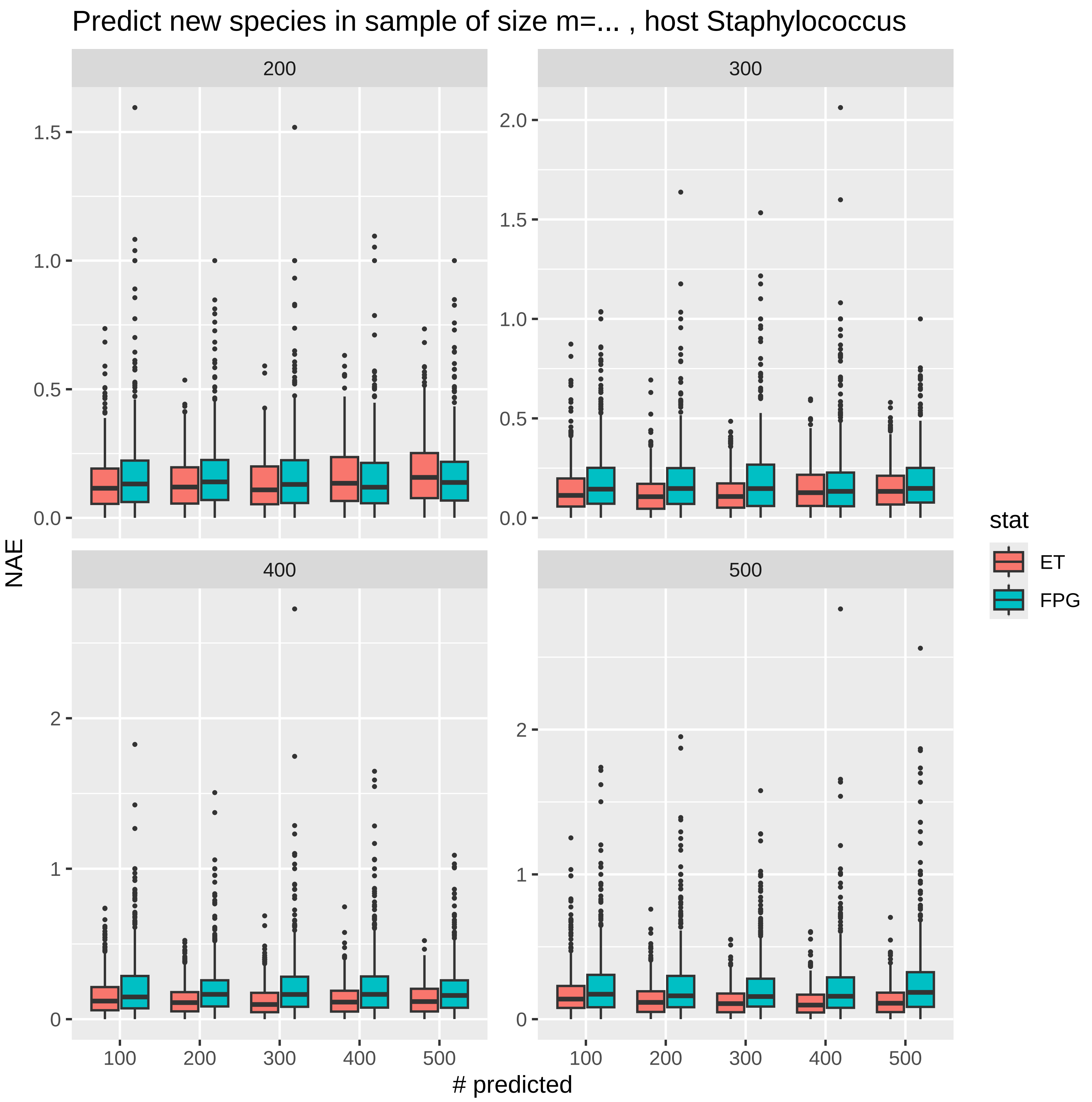

Predict new species in sample of size m=... , host Staphylococcus
200
300
400
500
NAE
### predicted
100
200
300
400
500
0.0
0.5
1.0
1.5
2.0
0
1
2
stat
ET
FPG

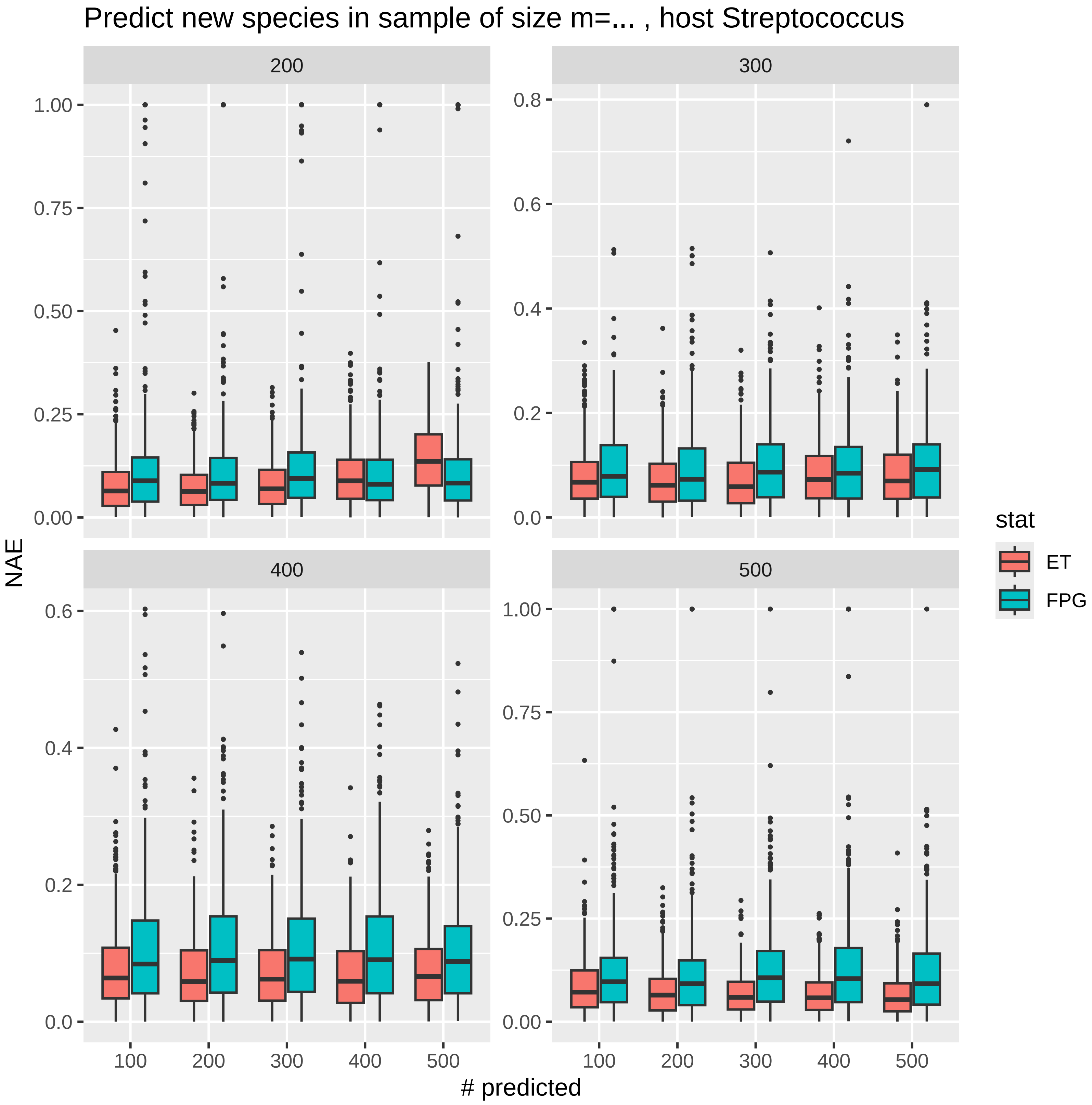
Predict new species in sample of size m=... , host Streptococcus
200
300
400
500
NAE
### predicted
100
200
300
400
500
1.00
0.75
0.50
0.25
0.00
0.8
0.6
0.4
0.2
0.0
stat
ET
FPG

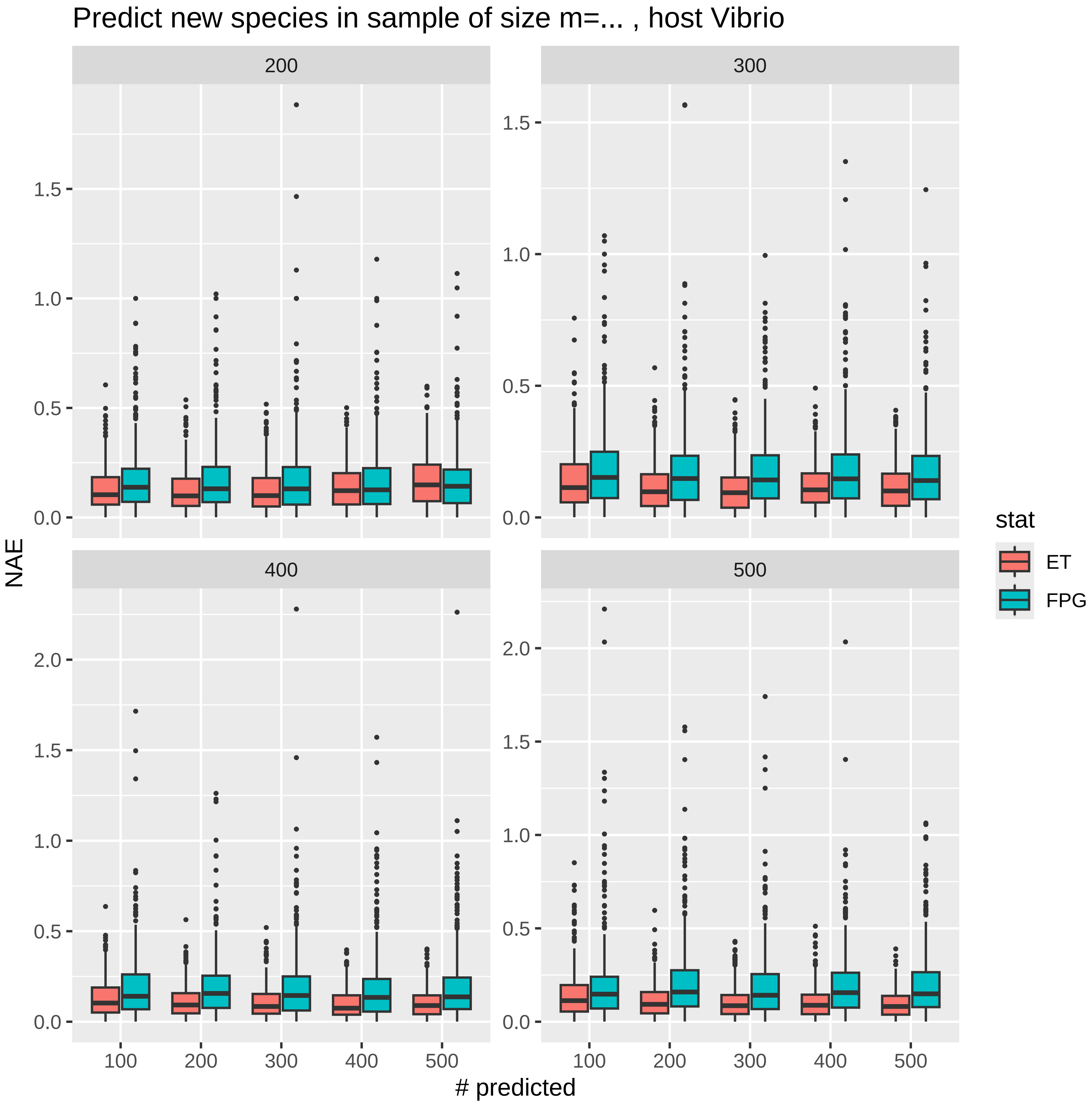
Predict new species in sample of size m=... , host Vibrio
200
300
400
500
NAE
### predicted
100
200
300
400
500
0.0
0.5
1.0
1.5
2.0
stat
ET
FPG

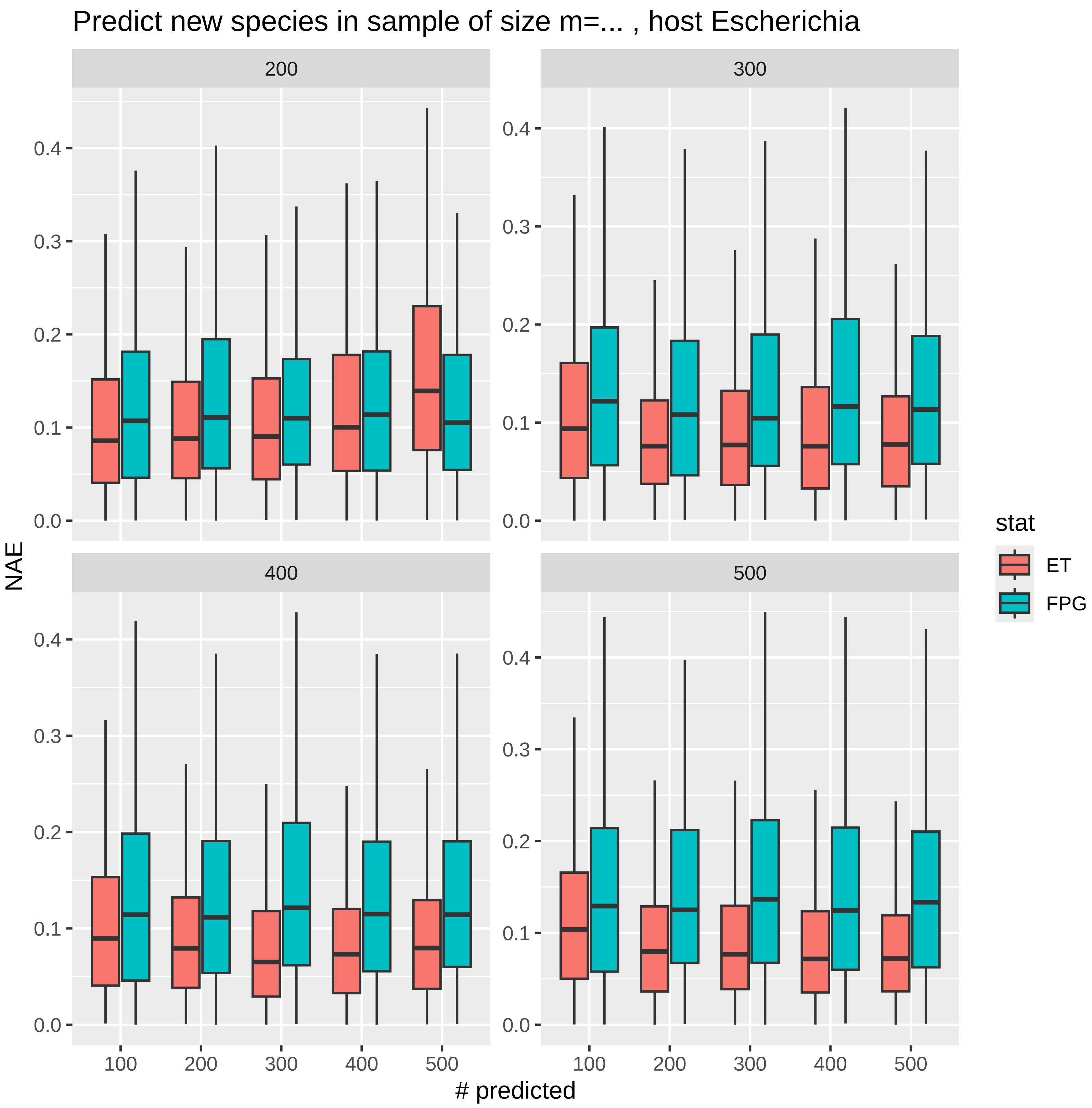

Predict new species in sample of size m=... , host Escherichia
200
300
400
500
NAE
### predicted
0.0
0.1
0.2
0.3
0.4
100
200
300
400
500
stat
ET
FPG

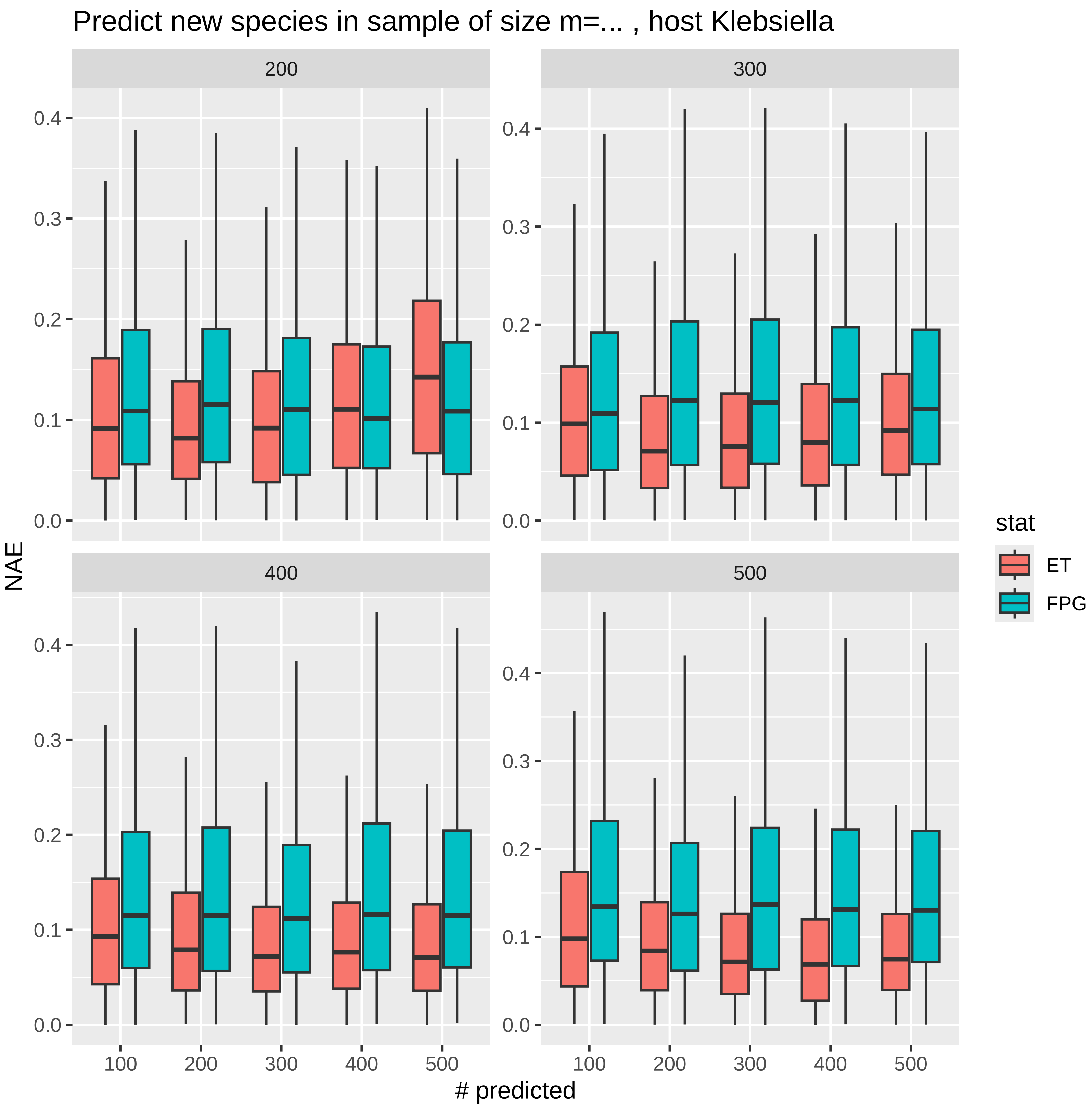

Predict new species in sample of size m=... , host Klebsiella
200
300
400
500
NAE
0.0
0.1
0.2
0.3
0.4
100
200
300
400
500
### predicted
stat
ET
FPG

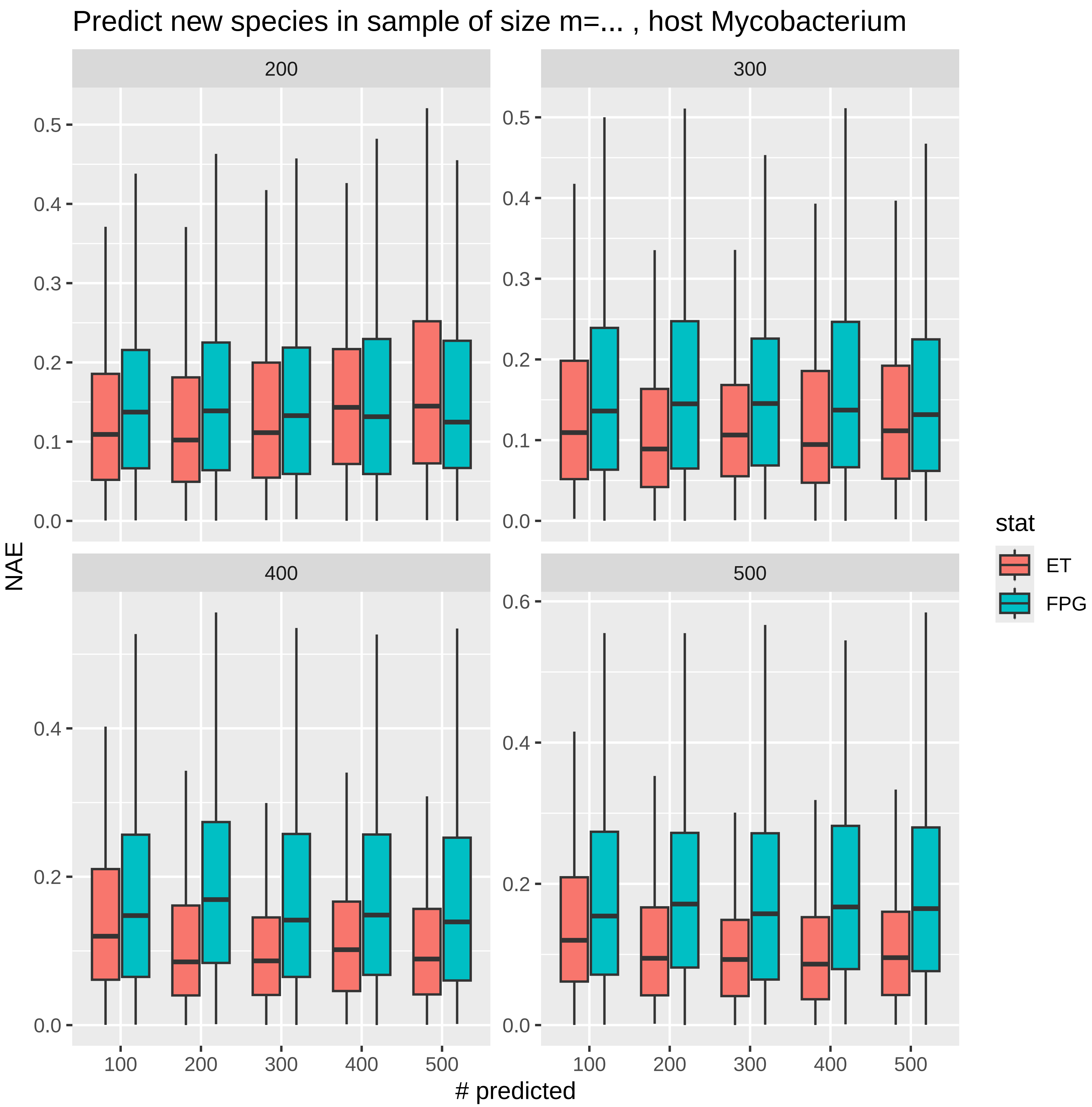

Predict new species in sample of size m=... , host Mycobacterium
200
300
400
500
NAE
### predicted
stat
ET
FPG
0.0
0.1
0.2
0.3
0.4
0.5
0.6
100
200
300
400
500

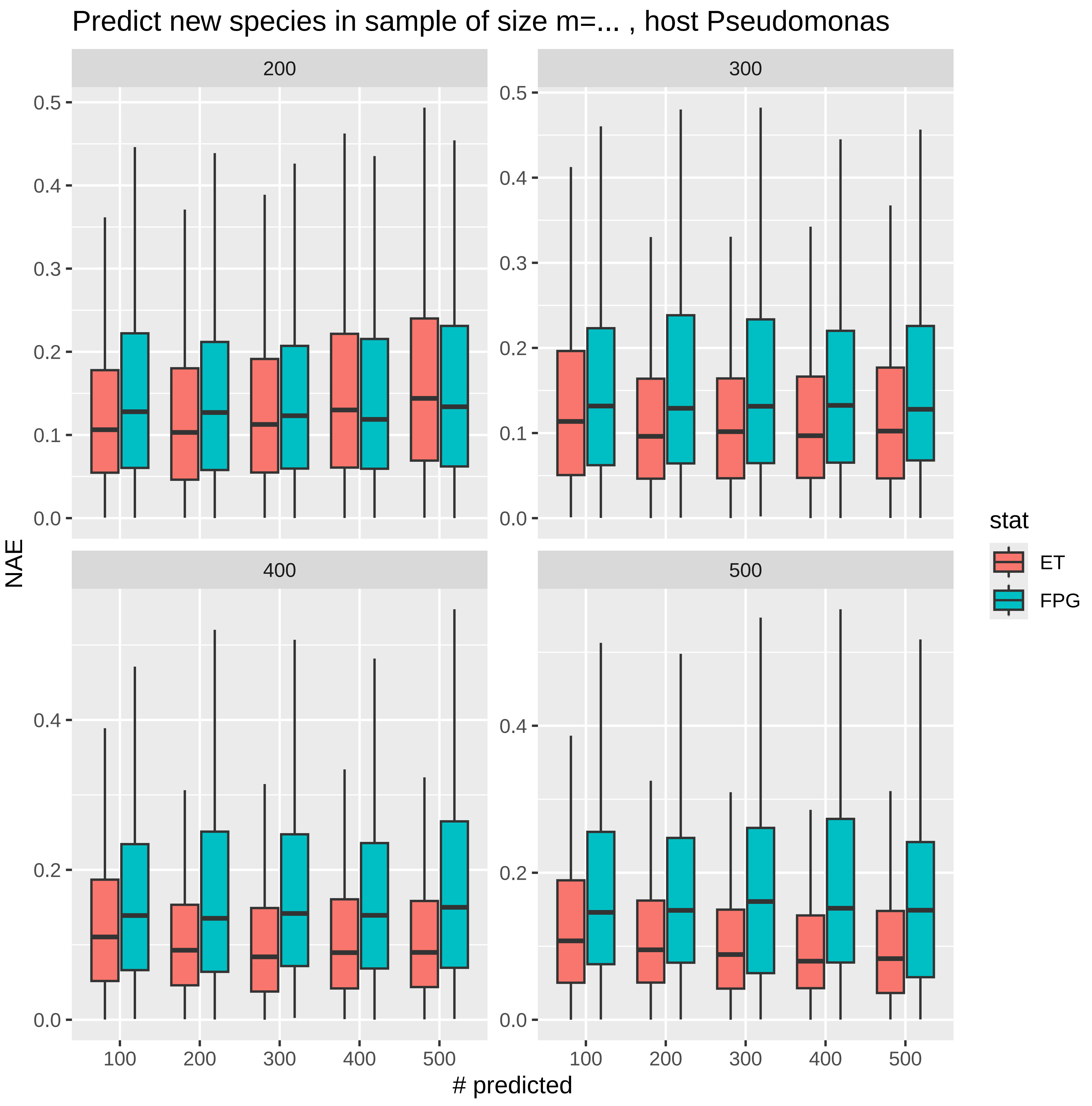
Predict new species in sample of size m=... , host Pseudomonas
200
300
400
500
0.5
0.4
0.3
0.2
0.1
0.0
NAE
100
200
300
400
500
### predicted
stat
ET
FPG

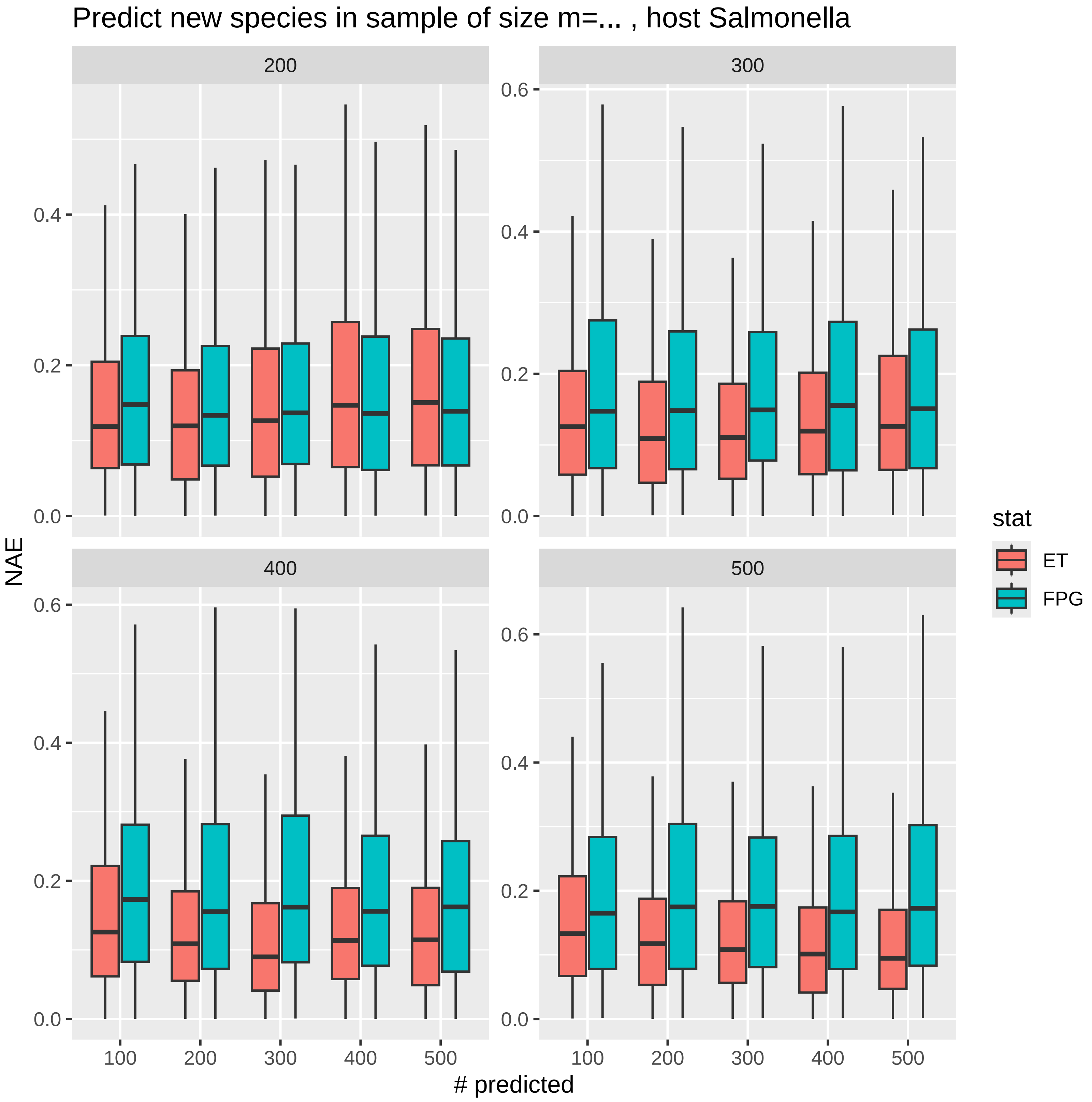

Predict new species in sample of size m=... , host Salmonella
200
300
400
500
NAE
### predicted
0.0
0.2
0.4
0.6
100
200
300
400
500
stat
ET
FPG

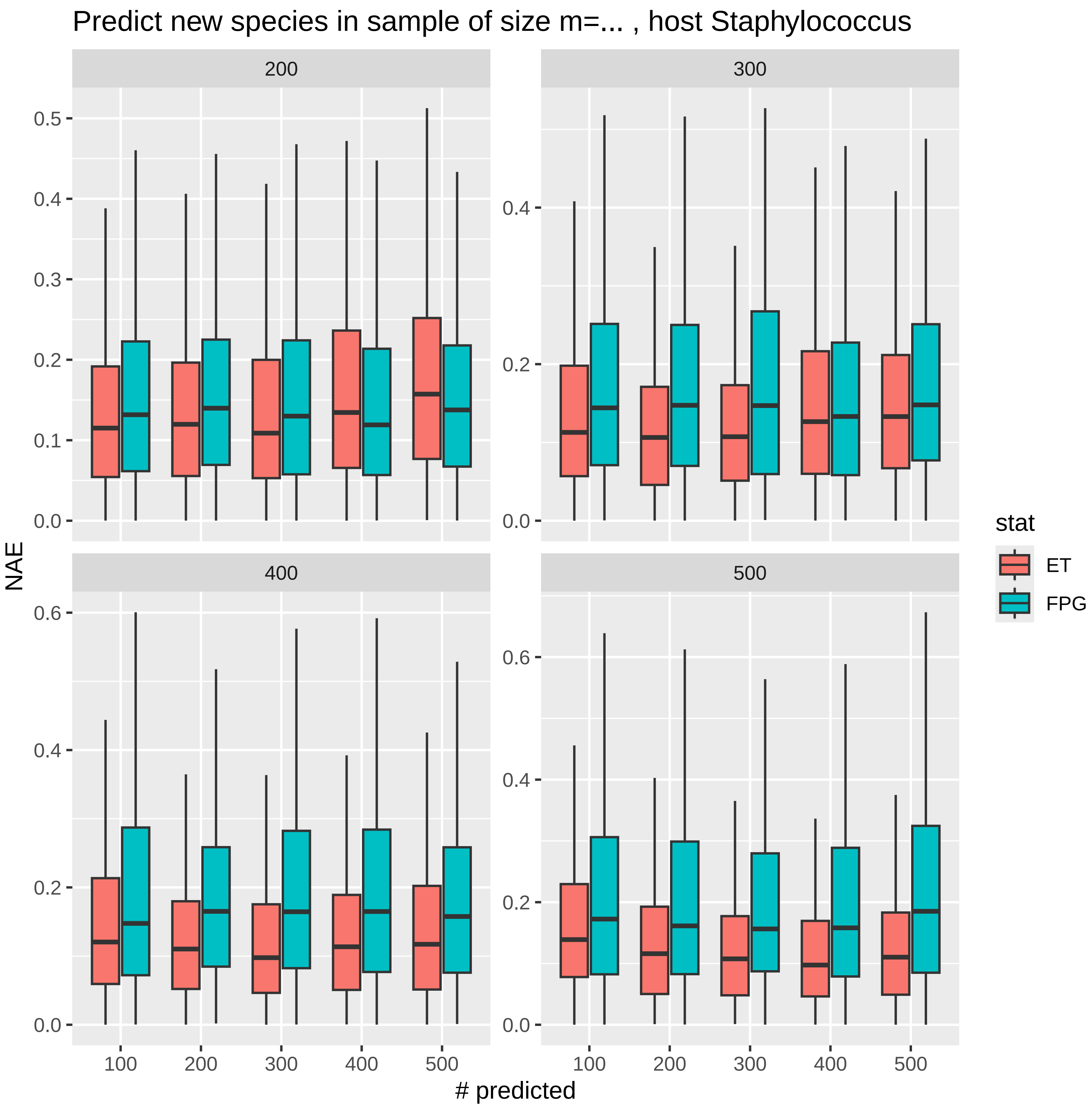

Predict new species in sample of size m=... , host Staphylococcus
200
300
400
500
0.5
0.4
0.3
0.2
0.1
0.0
0.6
NAE
### predicted
100
stat
ET
FPG

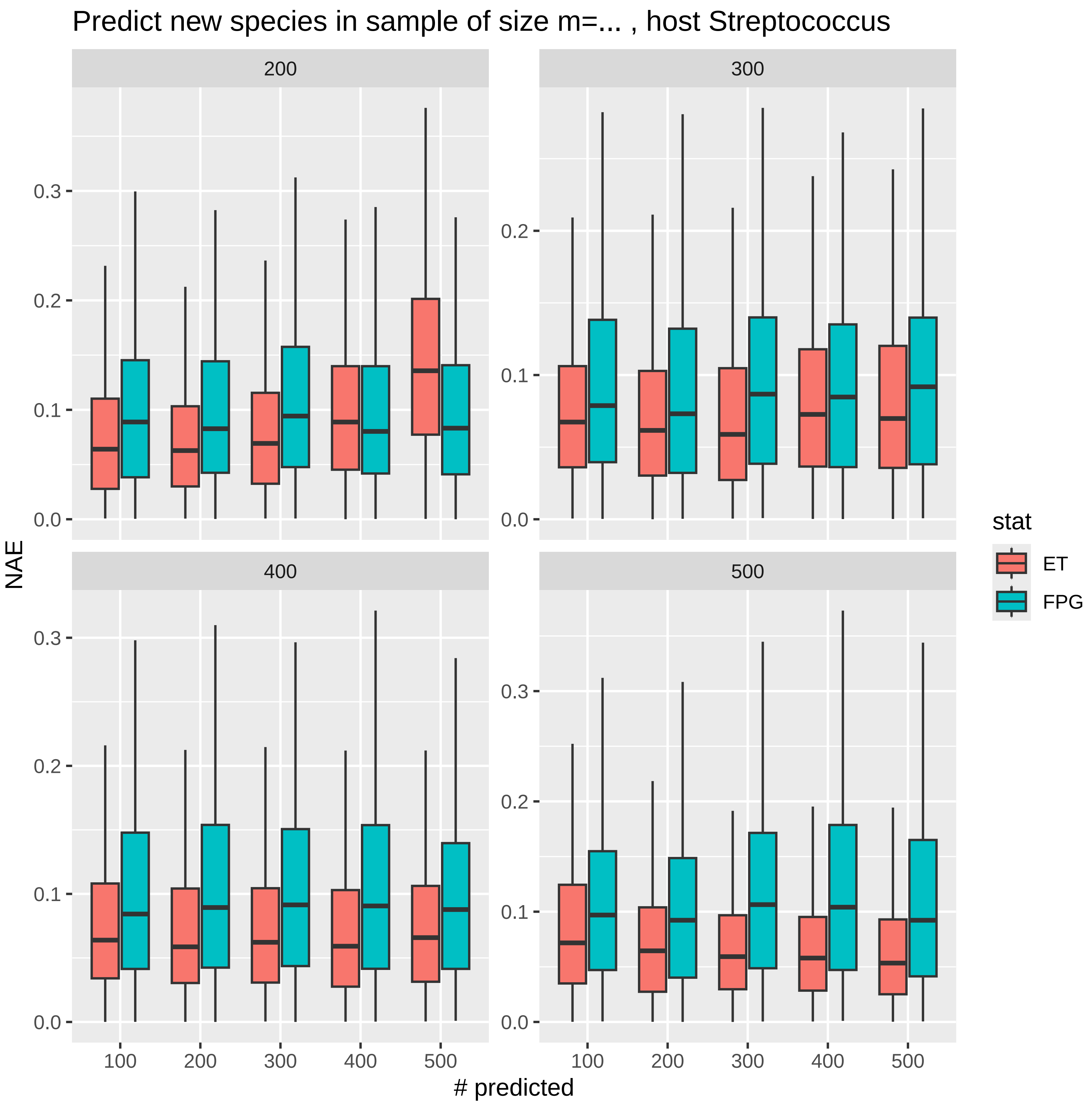
Predict new species in sample of size m=... , host Streptococcus
200
300
400
500
NAE
### predicted
100
200
300
400
500
0.0
0.1
0.2
0.3
stat
ET
FPG

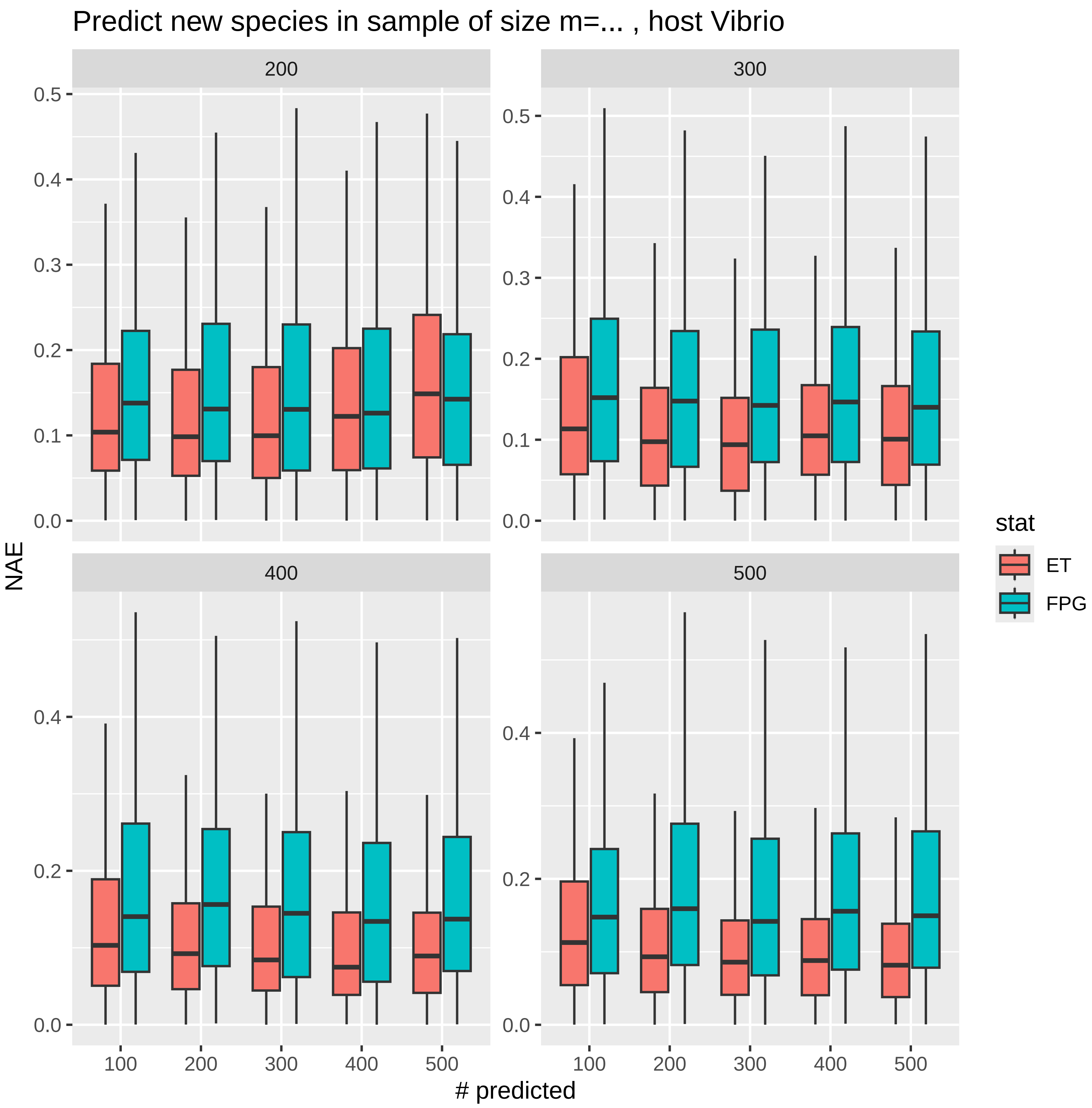

Predict new species in sample of size m=... , host Vibrio
200
300
400
500
0.0
0.1
0.2
0.3
0.4
0.5
NAE
100
200
300
400
500
### predicted
stat
ET
FPG

### 4 Exploring phage diversity using Hill numbers

#### 4.1 Diversity metrics - methods

Biological diversity describes how many species are present in a community and how evenly individuals are distributed among those species. A set of individuals of many rare species has a higher diversity than a set with only a few abundant species [2]. We quantify the biological diversity of phages in each host by means of Hill numbers ${}^{q}D(n)$ [3], defined as

$$ {}^{q}D(n) := \begin{cases} \exp\left(-\sum_{i=1}^{S_{obs}} \frac{a_i(n)}{n} \log \frac{a_i(n)}{n}\right), & \text{if } q = 1 \\ \left[\sum_{i}^{S_{obs}} \left(\frac{a_i(n)}{n}\right)^{q}\right]^{1/(1-q)}, & \text{otherwise} \end{cases} \tag{1} $$

which average the relative abundances of all species observed. These metrics depend on the order parameter $q > 0$, which determines its sensitivity to relative abundance, with increasing values of $q$ placing more weight on the most abundant species. Following standard conventions, we consider the case $q = 0$, where the Hill number effectively reduces to $S_{obs}$, the case $q = 1$, which is also referred to Hill-Shannon diversity, and the case $q = 2$, which yields the so-called Hill-Simpson diversity.

To allow for comparison between biodiversity measures in hosts with different sampling levels, we used an extrapolation method, i.e., given the initial $n$-individual sample, we estimated Hill numbers ${}^{q}\hat{D}(n+m)$ for an augmented sample of size $n + m$. For the estimation we followed the procedure of [4, 5] as implemented in the R package iNext [6].

#### 4.2 Phage diversity - results

Comparison of the phage diversity for the three orders of Hill numbers ($q = 0$, $q = 1$, and $q = 2$), for each different host, was performed based on extrapolated estimates from the 2024 dataset. All diversity curves reached the asymptote extrapolating to $10^4$, individual samples (Supplementary Figure 2). Regarding the mere number of phage species ($q = 0$), the *Escherichia* curve was the curve with the highest number (914.6), followed by *Vibrio*

(886.0), *Klebsiella* (848.7), *Streptococcus* (732.7), *Pseudomonas* (640.9), *Mycobacterium* (613.6), *Staphylococcus* (405.0), and *Salmonella* (392.1) (note that these are asymptotic values, while the true observed numbers $S_{obs}$ are in Table 1 in the main manuscript).

Considering the Hill-Shannon diversity ($q = 1$), the ranking a similar pattern to $q = 0$, with the highest asymptotic values for *Vibrio* (392.8), followed by *Klebsiella* (318.0), *Escherichia* (300.9), *Pseudomonas* (214.7), *Mycobacterium* (195.0), *Salmonella* (161.7), and *Staphylococcus* (150.8) – see also Figure 9.

Finally, the highest Hill-Simpson diversity (which is based on the dominant species and is obtained for $q = 2$) was in *Streptococcus* (239.2), followed at substantial distance by *Vibrio* (95.5), *Klebsiella* (76.5), *Escherichia* (69.1), *Pseudomonas* (56.8), *Salmonella* (53.), *Mycobacterium* (51.5), and *Staphylococcus* (36.9).

Extrapolation of the 2024 diversity estimates to the actual number of phages sampled in the 2025 allows a comparison between the two datasets. For $q = 0$, extrapolation of the Hill numbers is the Chao-Jost estimator from the main manuscript. The results for $q = 1$ are illustrated in Figure 9. The 2025 observed Hill-Shannon diversity numbers (solid circle markers) fall within, or very close to, the 95% confidence levels of the estimates extrapolated from the 2024 data (shaded areas) for six hosts (i.e., *Klebsiella*, *Escherichia*, *Mycobacterium*, *Pseudomonas*, *Salmonella*, and *Staphylococcus*). In contrast, the observed 2025 Hill-Shannon numbers were substantially above and below the estimated ranges for *Streptococcus* and *Vibrio* (the two hosts with the most diverse phage communities), respectively. The values of the Hill numbers for the 2024 and 2025 datasets, and the 2025 estimates from extrapolation of 2024 data, are reported in Table 1.

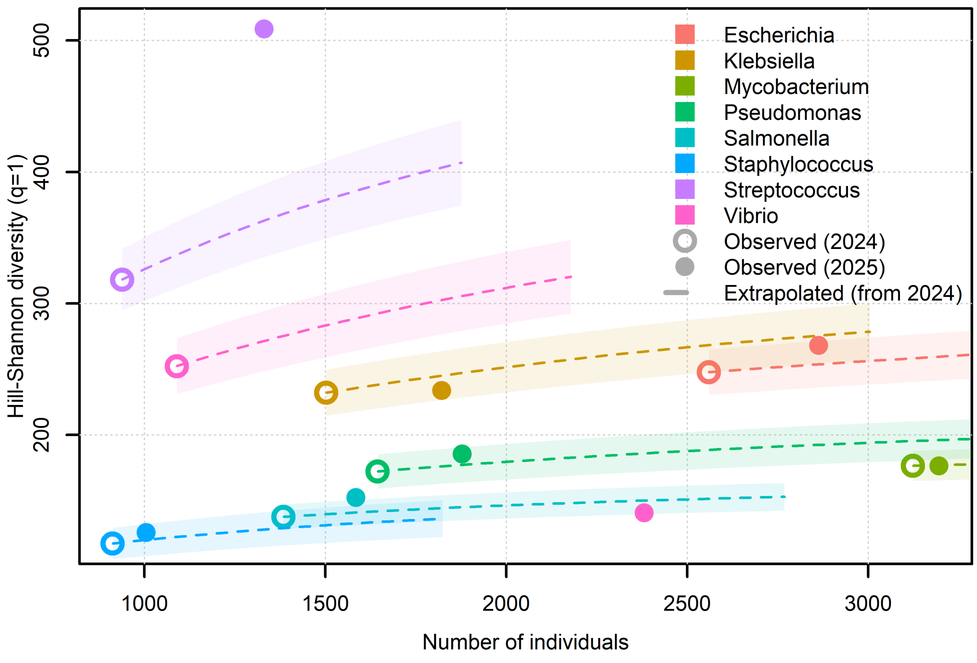


Figure 9: Shannon-Hill diversity of phages per bacterial host. The dashed lines are curves extrapolated by sampling from the 2024 data set, with 95% confidence intervals (shaded areas). Circle and solid circle makers represent the observed diversities in 2024 and 2025, respectively. Diversity extrapolation from 2024 does not capture diversity measures from 2025 for genera *Vibrio* and *Streptococcus*.

| Host | 2024 | | 2025 (Extrap.) | | 2025 | |
|---|---|---|---|---|---|---|
| | $^0\hat{D}$ | 95% CI | $^0\hat{D}$ | 95% CI | $^0\hat{D}$ | 95% CI |
| *Escherichia* | 716 | 694.93-737.07 | 749.44 | 726.90-771.98 | 826 | 801.97-850.03 |
| *Klebsiella* | 532 | 508.61-555.39 | 587.81 | 558.86-616.76 | 616 | 588.08-643.92 |
| *Mycobacterium* | 563 | 546.47-579.53 | 566.28 | 549.49-583.06 | 574 | 558.23-589.77 |
| *Pseudomonas* | 448 | 428.12-467.88 | 476.90 | 454.66-499.15 | 517 | 495.21-538.79 |
| *Salmonella* | 332 | 316.40-347.60 | 346.51 | 329.11-363.90 | 370 | 355.89-384.11 |
| *Staphylococcus* | 279 | 263.73-294.27 | 291.67 | 275.24-308.09 | 313 | 298.61-327.39 |
| *Streptococcus* | 449 | 429.81-468.19 | 536.55 | 507.62-565.48 | 705 | 682.10-727.90 |
| *Vibrio* | 476 | 455.32-496.68 | 711.10 | 662.44-759.75 | 602 | 576.41-627.59 |
| | $^1\hat{D}$ | 95% CI | $^1\hat{D}$ | 95% CI | $^1\hat{D}$ | 95% CI |
| *Escherichia* | 247.72 | 231.04-264.39 | 253.66 | 236.58-270.74 | 268.01 | 251.55-284.47 |
| *Klebsiella* | 232.01 | 215.17-248.86 | 245.04 | 226.83-263.24 | 233.70 | 212.98-254.41 |
| *Mycobacterium* | 176.47 | 165.95-186.98 | 176.96 | 166.42-187.50 | 176.25 | 166.61-185.90 |
| *Pseudomonas* | 172.20 | 159.80-184.59 | 177.13 | 164.28-189.98 | 185.33 | 173.12-197.55 |
| *Salmonella* | 137.68 | 126.75-148.61 | 140.89 | 129.70-152.08 | 152.25 | 141.88-162.61 |
| *Staphylococcus* | 117.32 | 106.20-128.45 | 120.04 | 108.61-131.47 | 125.70 | 114.13-137.27 |
| *Streptococcus* | 318.06 | 297.69-338.43 | 363.04 | 338.72-387.36 | 508.68 | 476.75-540.61 |
| *Vibrio* | 252.24 | 231.86-272.62 | 328.59 | 300.04-357.15 | 140.55 | 130.63-150.46 |
| | $^2\hat{D}$ | 95% CI | $^2\hat{D}$ | 95% CI | $^2\hat{D}$ | 95% CI |
| *Escherichia* | 67.71 | 59.83-75.59 | 67.90 | 59.97-75.83 | 66.49 | 59.97-73.02 |
| *Klebsiella* | 73.38 | 63.56-83.19 | 74.00 | 64.02-83.98 | 61.67 | 53.97-69.36 |
| *Mycobacterium* | 50.94 | 46.48-55.40 | 50.96 | 46.50-55.42 | 50.15 | 45.96-54.34 |
| *Pseudomonas* | 55.23 | 47.77-62.68 | 55.46 | 47.94-62.97 | 55.83 | 49.30-62.35 |
| *Salmonella* | 52.17 | 45.17-59.16 | 52.41 | 45.35-59.47 | 58.13 | 51.02-65.24 |
| *Staphylococcus* | 35.58 | 29.32-41.85 | 35.71 | 29.40-42.02 | 35.32 | 29.42-41.23 |
| *Streptococcus* | 194.48 | 171.27-217.70 | 207.11 | 181.14-233.09 | 312.72 | 277.97-347.47 |
| *Vibrio* | 88.65 | 72.95-104.36 | 92.71 | 75.61-109.80 | 34.21 | 30.51-37.92 |

Table 1: Hill numbers and their 95% confidence intervals (CIs) obtained from the 2024 data (left columns), extrapolated from the 2024 dataset to 2025 sampling (mid columns), and obtained from the 2025 dataset (right columns). $^0\hat{D}$ is Hill number of order zero, corresponding to the number of species, $^1\hat{D}$ is Hill-Shannon diversity, and $^2\hat{D}$ is Hill-Simpson diversity.

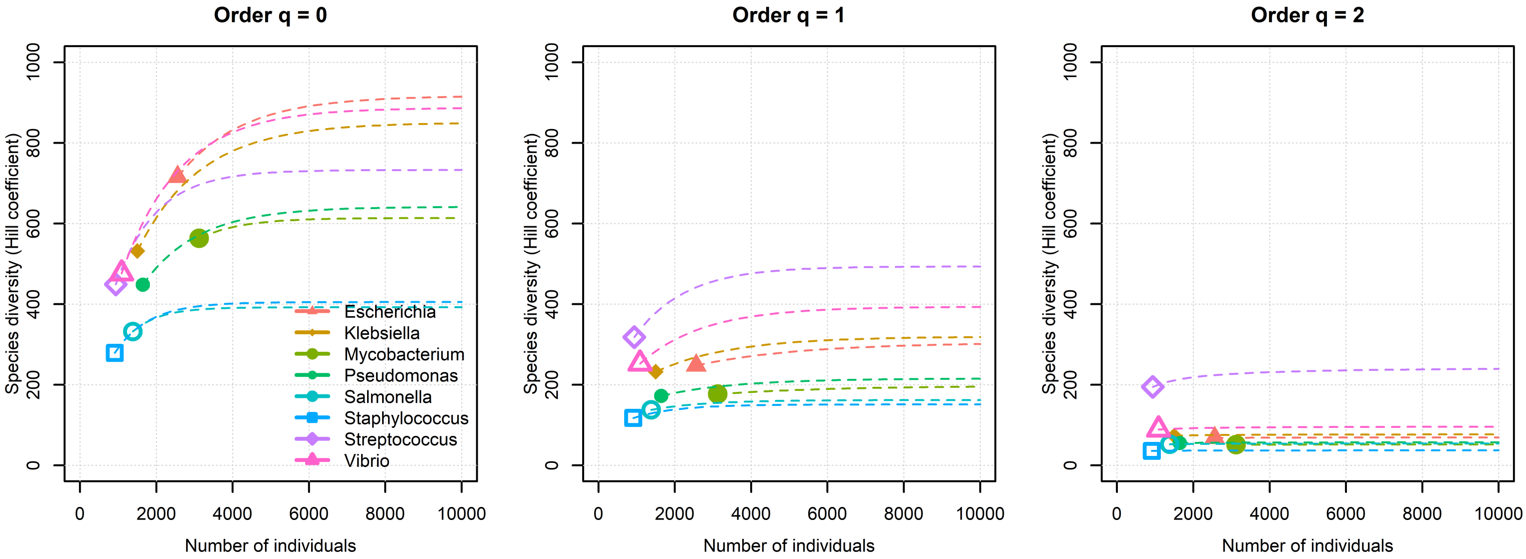


Figure 10: Extrapolating the Hill diversity curves from the 2024 data shows asymptotic convergence as sampling effort increases.

[2] M. Roswell, J. Dushoff, R. Winfree, A conceptual guide to measuring species diversity, Oikos 130 (3) 321–338. doi:10.1111/oik.07202.

[3] M. O. Hill, Diversity and evenness: A unifying notation and its consequences 54 (2) 427–432. doi:10.2307/1934352.

[4] A. Chao, N. J. Gotelli, T. C. Hsieh, E. L. Sander, K. H. Ma, R. K. Colwell, A. M. Ellison, Rarefaction and extrapolation with Hill numbers: a framework for sampling and estimation in species diversity studies, Ecological Monographs 84 (1) 45–67. doi:10.1890/13-0133.1.

[5] T.-J. Shen, A. Chao, C.-F. Lin, Predicting the number of new species in further taxonomic sampling, Ecology 84 (3) 798–804. doi:10.1890/0012-9658(2003)084[0798:PTNONS]2.0.CO;2.

[6] T. C. Hsieh, K. H. Ma, A. Chao, iNEXT: an R package for rarefaction and extrapolation of species diversity (Hill numbers), Methods in Ecology and Evolution 7 (12) 1451–1456. doi:10.1111/2041-210X.12613.
